# Mitochondrial calcium signaling mediated transcriptional regulation of keratin filaments is a critical determinant of melanogenesis

**DOI:** 10.1101/2023.05.26.542250

**Authors:** Jyoti Tanwar, Kriti Ahuja, Akshay Sharma, Paras Sehgal, Gyan Ranjan, Farina Sultan, Anshu Priya, Manigandan Venkatesan, Vamsi K Yenamandra, Archana Singh, Muniswamy Madesh, Sridhar Sivasubbu, Rajender K Motiani

**Author notes:** Corresponding author Address for correspondence: Dr. Rajender K Motiani Assistant Professor Regional Centre for Biotechnology (RCB) Faridabad, Delhi-NCR, India.

## Abstract

Mitochondria are versatile organelles that regulate several physiological functions. Many mitochondria-controlled processes are driven by mitochondrial Ca^2+^ signaling. However, role of mitochondrial Ca^2+^ signaling in melanosome biology remains unknown. Here, we show that pigmentation requires mitochondrial Ca^2+^ uptake. *In vitro* gain and loss of function studies demonstrated that Mitochondrial Ca^2+^ Uniporter (MCU) is crucial for melanogenesis while the MCU rheostats, MCUb and MICU1 negatively control melanogenesis. Zebrafish and mouse models showed that MCU plays a vital role in pigmentation *in vivo*. Mechanistically, MCU controls activation of transcription factor NFAT2 to induce expression of three keratins (keratin 5, 7 and 8), which we report as positive regulators of melanogenesis. Interestingly, keratin 5 in turn modulates mitochondrial Ca^2+^ uptake thereby this signaling module acts as a negative feedback loop that fine-tunes both mitochondrial Ca^2+^ signaling and melanogenesis. Mitoxantrone, an FDA approved drug that inhibits MCU, decreases physiological melanogenesis. Collectively, our data demonstrates a critical role for mitochondrial Ca^2+^ signaling in vertebrate pigmentation and reveal the therapeutic potential of targeting MCU for clinical management of pigmentary disorders. Given the centrality of mitochondrial Ca^2+^ signaling and keratin filaments in cellular physiology, this feedback loop may be functional in a variety of other pathophysiological conditions.

**Highlights:**

- MCU complex mediated mitochondrial Ca^2+^ uptake is a novel regulator of vertebrate pigmentation
- Keratin filaments bridge mitochondrial Ca^2+^ signaling to melanosome biogenesis and maturation
- Transcription factor NFAT2 connects mitochondrial Ca^2+^ dynamics to keratins expression
- MCU-NFAT2-Keratin 5 signaling module generates a negative feedback loop to maintain mitochondrial Ca^2+^ homeostasis and to ensure optimal melanogenesis
- Inhibiting MCU with mitoxantrone, an FDA approved drug, leads to reduction in physiological pigmentation

## Introduction

Mitochondria are emerging as robust signaling organelle that modulate functions of other organelles and thereby regulate cellular physiology (Picard & Shirihai, 2022). This inter organellar crosstalk plays an important role in their functioning and adaptation to changes in the cellular environment (Gottschling & Nystrom, 2017; Murley & Nunnari, 2016). Mitochondria can sense extracellular as well as intracellular cues and in response generate output signals for tuning activity of other organelles. One of the key signaling pathways that regulates mitochondria mediated modulation of other organelle functioning and thus overall cellular physiology is mitochondrial Ca^2+^ dynamics (Granatiero, De Stefani et al., 2017; Pathak & Trebak, 2018; Picard & Shirihai, 2022).

Mitochondrial Ca^2+^ signaling plays a vital role in regulating diverse cellular processes (Granatiero et al., 2017). Ca^2+^ uptake into the mitochondrial matrix is mediated via a highly Ca^2+^ selective channel i.e. Mitochondrial Calcium Uniporter (MCU) (Baughman, Perocchi et al., 2011; De Stefani, Raffaello et al., 2011). The Uniporter complex consists of pore-forming subunit MCU along with regulatory proteins namely MItochondrial Ca^2+^ Uptake proteins (MICU1/2), MCU regulatory subunit b (MCUb), Essential MCU Regulator (EMRE) and Mitochondrial Ca^2+^ Uniporter Regulator 1 (MCUR1) (Kamer & Mootha, 2015; Pathak & Trebak, 2018; Tanwar, Singh et al., 2021). Wherein, MICU1/2 act as gatekeepers of MCU complex and allow mitochondrial Ca^2+^ influx upon rise in inner-mitochondrial space Ca^2+^ concentration (Pathak & Trebak, 2018; Tanwar et al., 2021). EMRE tethers and stabilizes MICU1/MICU2 to the MCU complex (Sancak, Markhard et al., 2013). While MCUb acts as a dominant negative form of channel pore forming unit (MCU) and thereby it negatively regulates mitochondrial matrix Ca^2+^ uptake (Pathak & Trebak, 2018; Tanwar et al., 2021). The MCU complex plays a critical role in regulating several cellular functions including bioenergetics, autophagy, cytosolic Ca^2+^ buffering, secretory functions, cell proliferation/migration and cell survival (Carvalho, Stathopulos et al., 2020; Giorgi, Marchi et al., 2018). However, the significance of mitochondrial Ca^2+^ homeostasis and functional relevance of mitochondrial Ca^2+^ handling proteins in pigmentation biology remains undetermined.

Pigmentation is a complex physiological phenomenon that protects skin from UV induced DNA damage (Natarajan, Ganju et al., 2014a; Wasmeier, Hume et al., 2008). Inefficient pigmentation predisposes to skin cancers and perturbations in this pathway leads to pigmentary disorders such as vitiligo, melasma, Dowling Degos etc. (Natarajan et al., 2014a; Wasmeier et al., 2008). Pigmentation is an outcome of melanin synthesis (melanogenesis) in highly specialized lysosome-related organelles known as melanosomes. There are four stages of melanosomes wherein stage I-II are immature non-melanized while stage III-IV are highly melanized (Natarajan et al., 2014a; Wasmeier et al., 2008). The role of mitochondria in regulating melanosome biology and thereby pigmentation has just started to emerge (Rosania, 2005). Interestingly, mitochondria are reported to make physical contacts with melanosome via mitofusin 2 (MFN2) (Daniele, Hurbain et al., 2014). We recently demonstrated that MFN2 negatively regulates melanogenesis by modulating mitochondrial ROS generation (Tanwar, Saurav et al., 2022a). Similarly, mitochondria localized K^+^- dependent Na^+^/Ca^2+^ exchanger 5 (NCKX5), also known as SLC24A5, regulates melanosome biogenesis and melanin synthesis (Zhang, Gong et al., 2019). Nonetheless, role of mitochondrial Ca^2+^ dynamics in melanosome biology remains completely unknown.

Here, we demonstrate a critical role of mitochondrial Ca^2+^ signaling in pigmentation. Using two independent *in vitro* (mouse B16 cells and primary human melanocytes) and two distinct *in vivo* (zebrafish and knockin mice) models, we show that mitochondrial Ca^2+^ levels positively regulates melanogenesis. Notably, increase in melanogenesis is directly associated with mitochondrial Ca^2+^ levels. Mechanistically, alterations in mitochondrial Ca^2+^ dynamics lead to activation and nuclear translocation of transcription factor NFAT2. NFAT2 in turn regulates transcription of keratin 5, 7 and 8. These keratins enhance melanogenesis by augmenting melanosome biogenesis and maturation. Further, keratin 5 modulates mitochondrial Ca^2+^ uptake thereby this signaling cascade functions as a feedback loop to ensure optimum melanogenesis and to maintain mitochondrial Ca^2+^ homeostasis.

Interestingly, mutations in keratin 5 are associated with human pigmentary disorders such as Dowling Degos and epidermolysis bullosa with mottled pigmentation. Finally, inhibition of MCU with an FDA approved drug mitoxantrone decreases physiological melanogenesis thereby highlighting potential of targeting mitochondrial Ca^2+^ dynamics for managing pigmentary disorders.

## Results

### Pigmentation is associated with enhanced mitochondrial Ca^2+^ uptake

The functional significance of mitochondrial Ca^2+^ signaling in pigmentation is not studied yet. Therefore, we started by examining the mitochondrial Ca^2+^ dynamics during pigmentation. First of all, we measured mitochondrial Ca^2+^ uptake in B16 mouse melanoma cells while they were synchronously pigmenting in a low density (LD) culturing model. During LD pigmentation, non-pigmented B16 cells are seeded at a density of 100 cells/cm^2^ and over a period of 6-8 days these cells become highly pigmented (Motiani, Tanwar et al., 2018; Natarajan, Ganju et al., 2014b) (**Figure 1A**). The LD pigmentation model closely recapitulates human melanogenic pathways (Motiani et al., 2018; Natarajan et al., 2014b; Sultan, Basu et al., 2022; Tanwar, Sharma et al., 2022b). We quantitated the increase in the melanogenesis during LD pigmentation model by performing melanin content assay, which showed a gradual increase in melanin synthesis from LD day 0 (D0) to LD day 4 (D4), LD day 5 (D5) and LD day 6 (D6) (**Figure 1B**). We used genetically encoded Calcium measuring organelle-Entrapped Protein Indicators (CEPIA) for studying mitochondrial Ca^2+^ dynamics. CEPIA serves as a powerful tool for measuring intraorganellar Ca^2+^ dynamics (Suzuki, Kanemaru et al., 2014). We transfected B16 cells with a mitochondrial matrix specific CEPIA probe (CEPIA2mt) and temporally analyzed mitochondrial Ca^2+^ uptake during LD pigmentation model. We used histamine, a physiological agonist, for studying mitochondrial Ca^2+^ uptake. Histamine releases Ca^2+^ from the endoplasmic reticulum via Ca^2+^ release channel inositol-1,4,5-trisphosphate receptors (IP3Rs) thereby enhancing cytosolic Ca^2+^ levels. This rise in cytosolic Ca^2+^ stimulates mitochondrial Ca^2+^ uptake via

**Figure 1.**
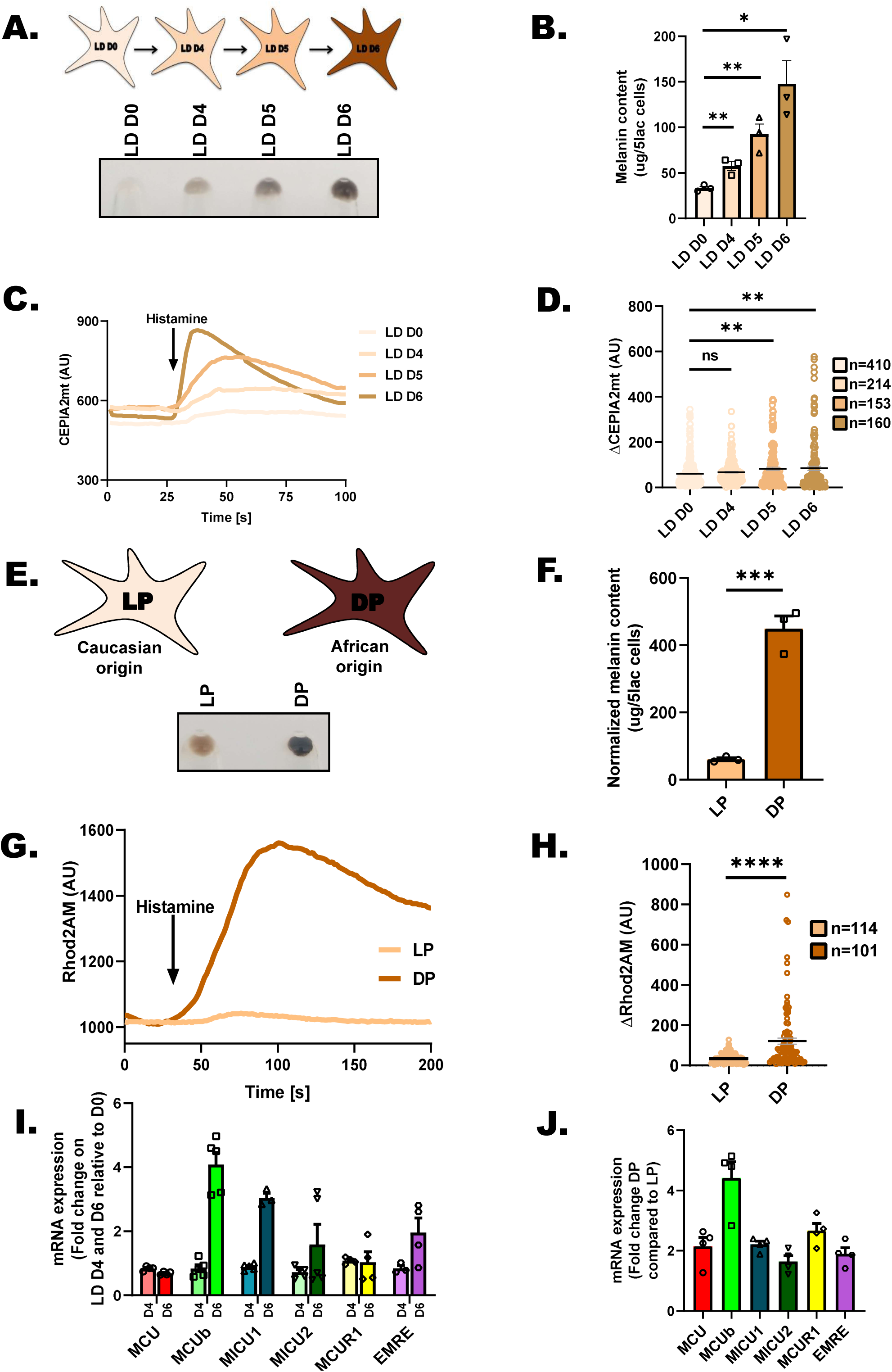
Mitochondrial Ca^2+^ uptake is positively associated with melanogenesis. **(A)** Representative B16 cell pellet pictures of LD day 0, LD day 4, LD day 5, and LD day 6 (N=3). **(B)** Melanin content estimation of B16 cells on LD day 0, LD day 4, LD day 5, and LD day 6 (N=3). **(C)** Representative mitochondrial Ca^2+^ imaging traces of CEPIA2mt on LD day 0, LD day 4, LD day 5, and LD day 6 B16 cells stimulated with 100µM histamine. **(D)** Quantitation of mitochondrial Ca^2+^ uptake by calculating ΔCEPIA2mt on LD day 0, LD day 4, LD day 5, and LD day 6 B16 cells stimulated with 100µM histamine where “n” denotes the number of ROIs. **(E)** Representative pellet pictures of Lightly pigmented (LP) and Darkly pigmented (DP) primary human melanocytes (N=3). **(F)** Melanin content estimation of LP and DP primary human melanocytes (N=3). **(G)** Representative mitochondrial Ca^2+^ imaging traces of LP and DP primary human melanocytes stimulated with 100µM histamine. **(H)** Quantitation of ΔRhod-2 in LP and DP primary human melanocytes stimulated with 100µM histamine where “n” denotes the number of ROIs. **(I)** qRT–PCR analysis showing relative mRNA expression of MCU complex components (MCU, MCUb, MICU1, MICU2, MCUR1 and EMRE) in B16 LD model on LD day 4 and LD day 6 (N=3-5). **(J)** qRT–PCR analysis showing relative mRNA expression of MCU complex components (MCU, MCUb, MICU1, MICU2, MCUR1 and EMRE) DP primary human melanocytes in comparison to LP primary human melanocytes (N=4). Data presented are mean ± S.E.M. For statistical analysis, unpaired student’s *t*-test was performed for panels B, F, H while one-way ANOVA followed by Tukey’s post-hoc test was performed for panel D using GraphPad Prism software. Here, NS means non-significant; * *p* <0.05; ** *p* < 0.01; *** *p* < 0.001 and **** *p* < 0.0001.

Mitochondrial Ca^2+^ Uniporter (MCU) complex. Our live cell mitochondrial Ca^2+^ imaging assays demonstrate that with increase in pigmentation, the mitochondrial Ca^2+^ uptake is enhanced (**Figure 1C and D**). This data suggests that during pigmentation there is an augmentation of mitochondrial Ca^2+^ dynamics.

We next investigated mitochondrial Ca^2+^ levels in primary human melanocytes with varying pigment levels i.e. lightly pigmented (LP) melanocytes of Caucasian origin and darkly pigmented (DP) melanocytes of Afro-American origin (Tanwar et al., 2022a) (**Figure 1E**). We quantitated the differential pigmentation in LP and DP primary human melanocytes by performing melanin content assay, which showed robust increase in melanin synthesis in DP as compared to LP (**Figure 1F**). We used mitochondrial Ca^2+^ measurement probe Rhod-2AM for examining differences in mitochondrial Ca^2+^ dynamics between primary LP and DP melanocytes. We studied histamine stimulated mitochondrial Ca^2+^ uptake in primary LP and DP melanocytes. Just like B16 LD pigmentation model, we observed that the mitochondrial Ca^2+^ uptake is higher in DP Afro-American melanocytes in comparison to LP Caucasian melanocytes (**Figure 1G and H**). Altogether, data from both B16 pigmentation model and primary human melanocytes highlight that increased pigmentation is associated with higher mitochondrial Ca^2+^ uptake.

Mitochondrial matrix Ca^2+^ uptake is a highly regulated process, which is mediated by Mitochondrial Ca^2+^ Uniporter (MCU) complex localized on inner mitochondrial membrane (Carvalho et al., 2020; Granatiero et al., 2017; Kamer & Mootha, 2015). The MCU complex consists of pore forming unit i.e. MCU; dominant negative isoform of MCU i.e. MCUb; gatekeepers MICU1/MICU2, MCUR1 and EMRE (Carvalho et al., 2020; Granatiero et al., 2017; Kamer & Mootha, 2015; Tanwar et al., 2021). Since we observed enhanced mitochondrial Ca^2+^ uptake, we evaluated the expression of MCU complex genes during pigmentation. We performed qRT-PCR for MCU, MCUb, MICU1, MICU2, MCUR1 and EMRE on LD day0, day4 & day6 samples and found that except MCUb all other MCU complex components remain largely unaltered during LD pigmentation (**Figure 1I**). Next, we carried out similar mRNA expression analysis of MCU complex components in primary LP and DP melanocytes. Interestingly, DP melanocytes have over three fold higher expression of MCUb while levels of other MCU complex components is increased by around two folds (**Figure 1J**). The expression data from B16 cells and primary melanocytes implicates that most likely enhanced mitochondrial Ca^2+^ uptake is not due to elevated expression of MCU complex components and could be associated with increased MCU activity. One of the most critical factors that regulates MCU activity is cytosolic Ca^2+^ levels (Finkel, Menazza et al., 2015; Rizzuto, Brini et al., 1993; Shanmughapriya, Rajan et al., 2015; Tanwar et al., 2021; Wang, Li et al., 2021). Increase in cytosolic Ca^2+^ concentration leads to higher Ca^2+^ in mitochondrial inter membrane space, which is sensed by MCU gatekeepers MICU1/2 eventually leading to opening of MCU pore and enhanced mitochondrial Ca^2+^ uptake. Interestingly, in our earlier study, we demonstrated that Store operated Ca^2+^ entry and resting cytosolic Ca^2+^ levels are higher in pigmented cells in comparison to non-pigmented cells (Motiani et al., 2018). The elevated cytosolic Ca^2+^ levels in pigmented cells could be the most likely driver of enhanced mitochondrial Ca^2+^ uptake in these cells. However, functional relevance of augmented mitochondrial Ca^2+^ levels in pigmentation biology remains completely obscure. Therefore, we investigated the role of MCU complex in pigmentation.

### MCU positively regulates melanogenesis

To understand the role of MCU in melanogenesis, we performed loss of function and gain of function studies in B16 cells. We first characterized MCU siRNA in B16 cells. Upon qRT– PCR analysis, we observed a drastic decrease in MCU mRNA levels in siMCU cells in comparison to control non-targeting siRNA (siNT) transfected B16 cells (**Supplementary Figure 1A**). We further validated the siRNA targeting MCU by performing western blotting. We observed a significant decrease (around 70%) in the MCU protein expression in the siMCU condition in comparison to siNT transfected cells (**Figure 2A**). The extent of decrease in MCU protein expression upon MCU knockdown was quantitated from several independent experiments (**Supplementary Figure 1B**). Next, we examined the functional outcome of MCU silencing on B16 mitochondrial Ca^2+^ dynamics. Using CEPIA2mt, we measured resting mitochondrial Ca^2+^ levels and histamine induced mitochondrial Ca^2+^ uptake upon MCU silencing. In these live cell imaging experiments, we observed that MCU silencing decreases basal/resting mitochondrial Ca^2+^ levels in B16 cells (**Figure 2B and 2C**). Further, we saw a significant decrease in mitochondrial Ca^2+^ uptake upon MCU silencing in comparison to siNT transfected control cells (**Figure 2B and 2D**). These results suggest that MCU silencing in B16 cells decreases both resting mitochondrial Ca^2+^ levels and agonist induced mitochondrial Ca^2+^ uptake.

**Figure 2.**
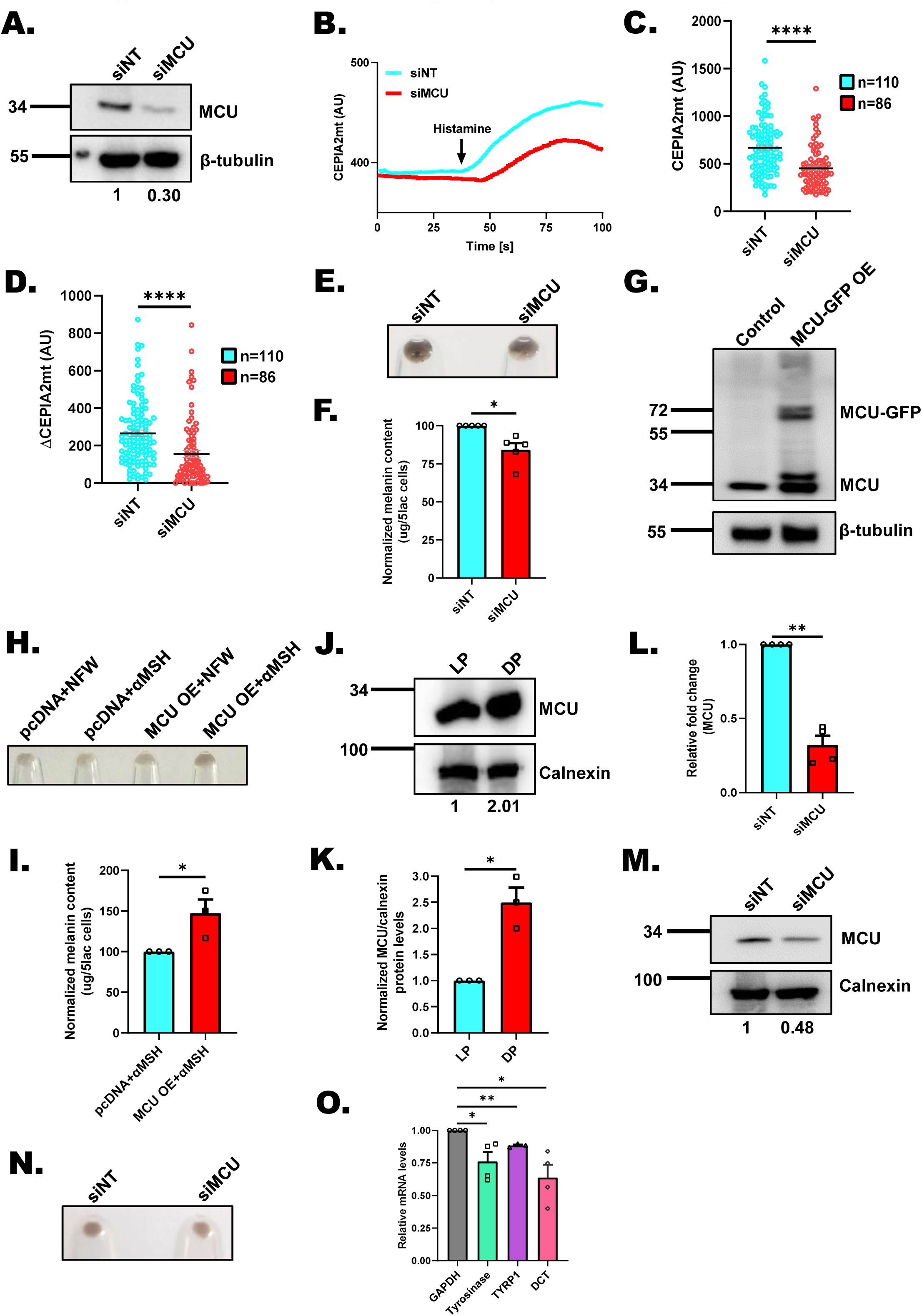
MCU positively regulates melanogenesis. **(A)** Representative western blot confirming siRNA based silencing of MCU on LD day 6 B16 cells. Densitometric analysis using ImageJ is presented below the blot (N=3). **(B)** Representative mitochondrial Ca^2+^ imaging traces of CEPIA2mt in siNon-Targeting (siNT) control and siMCU B16 cells stimulated with 100µM histamine. **(C)** Quantitation of resting mitochondrial Ca^2+^ with CEPIA2mt in siNT control and siMCU B16 cells where “n” denotes the number of ROIs. **(D)** Quantitation of mitochondrial Ca^2+^ uptake by calculating increase in CEPIA2mt signal (ΔCEPIA2mt) in siNT control and siMCU B16 cells stimulated with 100µM histamine where “n” denotes the number of ROIs. **(E)** Representative pellet pictures of siNT control and siMCU on LD day 6 (N=5). **(F)** Melanin content estimation of siNT and siMCU B16 cells on LD day 6 (N=5). **(G)** Representative western blot demonstrating ectopic expression of GFP-tagged human MCU in B16 cells (N=3). **(H)** Representative pellet pictures of pcDNA control plasmid and MCU-GFP overexpression either treated with nuclease free water (NFW) or αMSH (N=3). **(I)** Melanin content estimation of pcDNA control plasmid and MCU-GFP overexpression upon αMSH treatment (N=3). **(J)** Representative western blot showing an increase in MCU protein expression in DP primary human melanocytes in comparison to LP primary human melanocytes. Densitometric analysis using ImageJ is presented below the blot (N=3). **(K)** Densitometric quantitation showing increase in MCU protein levels in DP primary human melanocytes in comparison to LP primary human melanocytes (N=3). **(L)** qRT–PCR analysis showing decrease in MCU mRNA expression upon MCU silencing in primary human melanocytes (N=4). **(M)** Representative western blot confirming siRNA based silencing of MCU in primary human melanocytes. Densitometric analysis using ImageJ is presented below the blot (N=3). **(N)** Representative pellet pictures of siNT control and siMCU in primary human melanocytes (N=4). **(O)** qRT–PCR analysis showing decrease in Tyrosinase, TYRP1 and DCT mRNA expression upon MCU silencing in primary human melanocytes (N=4). Data presented are mean ± S.E.M. For statistical analysis, unpaired student’s *t*-test was performed for panels C, D while one sample *t*-test was performed for panels F, I, L, K, O using GraphPad Prism software. * *p* <0.05; ** *p* < 0.01 and **** *p* < 0.0001.

We then investigated the role of MCU in B16 LD melanogenesis. We performed siRNA mediated knockdown of MCU on LD day3 in B16 LD pigmentation model and examined LD cells on day6. The knockdown of MCU resulted in the visible decrease in LD day6 pigmentation as evident in the LD day6 pellet pictures (**Figure 2E**). We further quantitated the decrease in melanogenesis upon MCU silencing by performing melanin content assays. We observed that MCU silencing significantly decreased melanogenesis in B16 cells (**Figure 2F**). To further corroborate MCU’s role in melanogenesis we performed gain of function studies. For these experiments, we used non-pigmented B16 cells cultured in high density (HD) and induced melanogenesis in them with a physiological melanogenic stimuli α- melanocyte stimulating hormone (αMSH). We used αMSH as a stimuli for stimulating melanogenesis in B16 HD cells as it induces ER Ca^2+^ release and increases cytosolic Ca^2+^ levels and would thereby lead to mitochondrial Ca^2+^ uptake (Motiani et al., 2018). We overexpressed MCU in B16 cells by transfecting human MCU-GFP plasmid. We examined the MCU-GFP transfected B16 cells for GFP signal to gauge the transfection and observed over 60% transfection efficiency (**Supplementary Figure 1C**). Further, we confirmed MCU overexpression (OE) in B16 cells by performing western blotting. As MCU was tagged with GFP in the overexpression construct, we could observe a higher molecular-weight band corresponding to MCU-GFP in MCU-GFP overexpressing condition (**Figure 2G**). Next, we examined the effect of MCU overexpression on the αMSH-stimulated B16 pigmentation. Interestingly, we observed enhanced αMSH-induced pigmentation upon MCU overexpression in these cells. The pellet pictures from pcDNA (empty vector control)+NFW (nuclease free water; vehicle control for αMSH), pcDNA+αMSH, MCU-GFP OE+NFW and MCU-GFP OE+ αMSH conditions clearly showed increased pigmentation phenotype in MCU-GFP OE+αMSH condition (**Figure 2H**). We further quantitated these phenotypic changes by performing melanin-content assays and observed that indeed, the ectopic expression of MCU-GFP in B16 HD cells leads to a significant increase in αMSH-induced melanogenesis in comparison to empty vector control (**Figure 2I**). Taken together, our results demonstrate that MCU positively regulates melanogenesis in B16 cells.

In order to further substantiate the role of MCU in controlling melanogenesis, we next studied its impact on melanogenesis in primary human melanocytes. First of all, we examined MCU protein levels in darkly pigmented (DP) and lightly pigmented (LP) primary human melanocytes (**Figure 2J**). Our data shows that MCU levels are significantly higher (about 2 fold) in the darkly pigmented primary human melanocytes as compared to lightly pigmented primary human melanocytes (**Figure 2J and 2K**). This suggests that MCU expression is positively related to pigmentation levels in darkly pigmented primary human melanocyte cells. To examine the role of MCU in regulating melanogenesis in primary human melanocytes, we silenced MCU using human siRNAs in lightly pigmented primary human melanocytes. We used nucleofection technology for performing transfection in primary human melanocytes. We first validated MCU knockdown using qRT–PCR and western blot analysis. In qRT–PCR analysis, we observed a drastic decrease (around 70%) in MCU mRNA levels in siMCU condition in comparison to control siNT transfected primary human melanocytes (**Figure 2L**). Further, we observed about 50% decrease in the MCU protein expression in cells transfected with siMCU as compared to siNT transfected B16 cells (**Figure 2M**). The quantitation of the extent of decrease in MCU protein expression upon MCU knockdown was performed on several independent experiments and the quantitative data is presented in **Supplementary Figure 1D**. The knockdown of MCU resulted in visible decrease in melanogenesis as evident in the pellet pictures of siMCU in comparison to siNT control (**Figure 2N**). To further strengthen this phenotypic data, we analyzed mRNA expression of key melanogenic enzymes i.e. tyrosinase, tyrosinase-related protein 1 (TYRP1), and tyrosinase-related protein 2/Dopachrome Tautomerase (DCT) in the siNT and siMCU cells. We observed that MCU silencing in primary human melanocytes results in significant reduction in the expression of tyrosinase, TYRP1 and DCT (**Figure 2O**). Collectively, these experiments in three independent melanogenesis models (B16 LD model, B16 αMSH-induced pigmentation and melanogenesis in primary melanocytes) clearly demonstrate a critical role of MCU in regulating melanogenesis.

### MCUb and MICU1 negatively control melanogenesis

Next, we questioned whether the changes in melanogenesis upon altering MCU expression is due to concomitant fluctuations in mitochondrial Ca^2+^ levels or they are associated with some Ca^2+^ independent functions of MCU. To address this, we silenced MCUb, an important negative regulator of MCU (Raffaello, De Stefani et al., 2013) in B16 LD pigmentation model. We first characterized MCUb siRNA in B16 cells. Upon qRT–PCR analysis, we observed a significant reduction in MCUb mRNA levels in siMCUb treated cells in comparison to control siNT transfected B16 cells (**Figure 3A**). Next, we measured basal mitochondrial Ca^2+^ levels and histamine induced mitochondrial Ca^2+^ uptake upon MCUb silencing. We observed a significant increase in resting mitochondrial Ca^2+^ levels (**Figure 3B and 3C**) as well as mitochondrial Ca^2+^ uptake upon MCUb knockdown as compared to siNT control (**Figure 3B and D**). This data suggests that MCUb silencing enhances mitochondrial Ca^2+^ uptake. Then, we evaluated role of MCUb in B16 LD pigmentation model. We transfected B16 LD day3 cells with either siNT or siMCUb and assessed pigmentation phenotype on LD day6. We observed visible increase in LD day6 pigmentation as evident in the pellet pictures upon knockdown of MCUb as compared to control (**Figure 3E**). We further quantitated the increase in melanogenesis upon MCUb silencing by performing melanin content assays. We observed that MCUb silencing enhanced melanogenesis by ∼75% in B16 LD cells (**Figure 3F**). We next examined the expression of key melanosome structural protein (Pre-melanosome Protein 17 i.e. PMEL17 or GP100) and melanogenic enzymes (tyrosinase and DCT) upon the silencing of MCUb in B16 LD pigmentation model. We performed western blot analysis in order to evaluate the levels of GP100 and DCT in MCUb silenced B16 LD cells on LD day6. As presented in **Figure 3G**, siMCUb cells expressed significantly higher levels of GP100 and DCT as compared to siNT control cells on LD day 6. We then performed quantitation of the extent of increase in GP100 and DCT protein levels upon MCUb knockdown in several independent experiments (**Supplementary Figure 2A and B**). Further, we investigated the expression and activity of the rate-limiting enzyme in melanin synthesis pathway, i.e., tyrosinase. We performed tyrosinase protein analysis for its expression and DOPA (dopachrome generation) assay for its activity in siNT control and siMCUb cells on LD day6. Tyrosinase converts its substrate L-DOPA to Dopachrome that in turn yields a brown-to-black color on native gels. We observed that both the expression and activity of tyrosinase was significantly increased in siMCUb cells in comparison to siNT control cells (**Figure 3H**). We then quantitated the extent of the increase in both the expression and activity of tyrosinase upon MCUb knockdown in several independent experiments (**Supplementary Figure 2C and D**). This data suggests that MCUb is indeed a negative regulator of melanogenesis.

**Figure 3.**
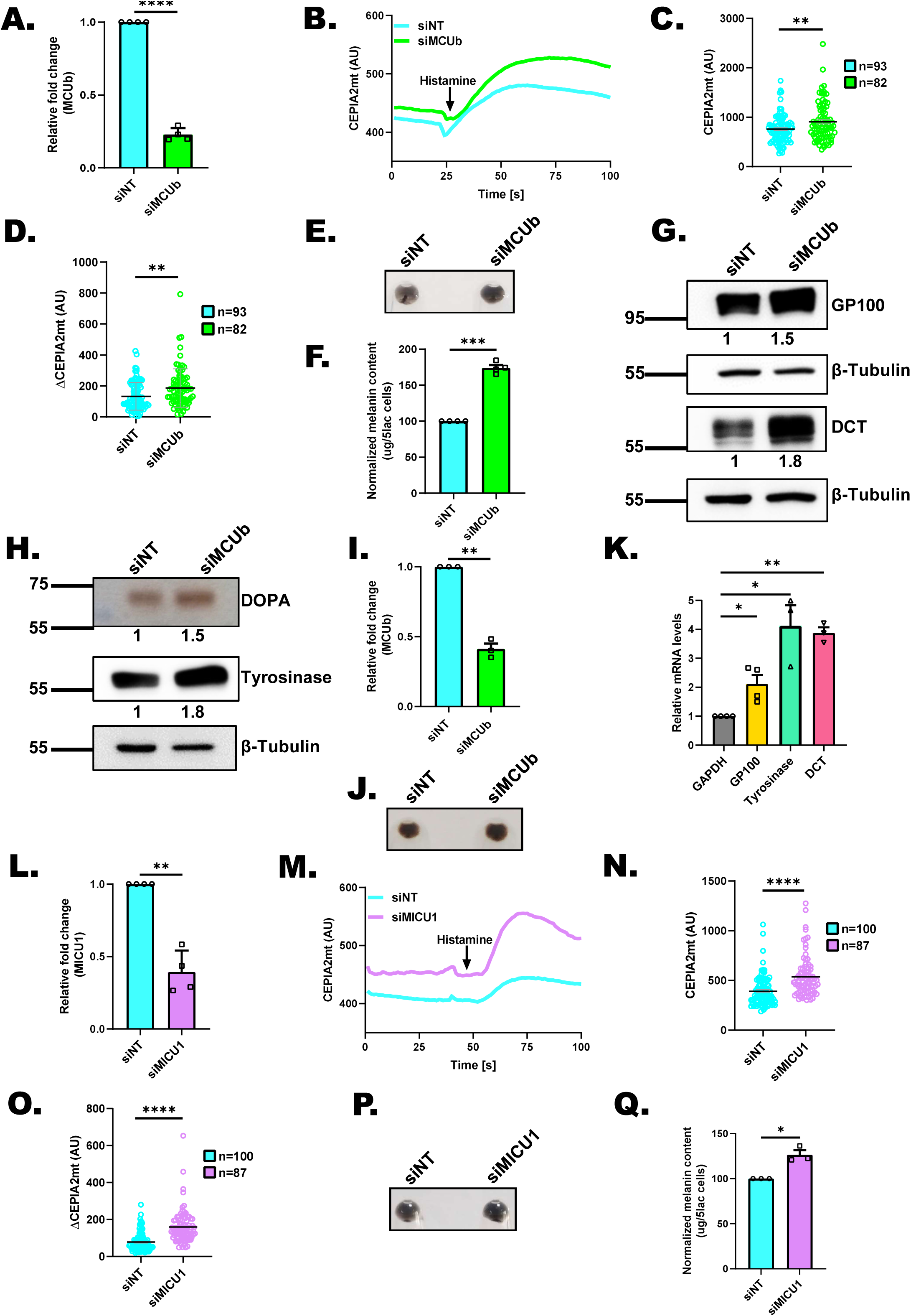
MCUb and MICU1 negatively control melanogenesis. **(A)** qRT–PCR analysis showing decrease in MCUb mRNA expression upon MCUb silencing in B16 cells (N=4). **(B)** Representative mitochondrial Ca^2+^ imaging traces of CEPIA2mt in siNT control and siMCUb B16 cells stimulated with 100µM histamine. **(C)** Quantitation of resting mitochondrial Ca^2+^ with CEPIA2mt in siNT control and siMCUb B16 cells where “n” denotes the number of ROIs. **(D)** Quantitation of mitochondrial Ca^2+^ uptake by calculating increase in CEPIA2mt signal (ΔCEPIA2mt) in siNT control and siMCUb B16 cells upon stimulation with 100µM histamine where “n” denotes the number of ROIs. **(E)** Representative pellet pictures of siNT control and siMCUb on LD day 6 (N=4). **(F)** Melanin content estimation in siNT and siMCUb B16 cells on LD day 6 (N=4). **(G)** Representative western blot showing expression of GP100 and DCT on LD day 6 upon MCUb silencing as compared to siNT control. Densitometric analysis using ImageJ is presented below the blot (N=3). **(H)** DOPA assay showing activity of tyrosinase enzyme (N=4) and representative western blot for tyrosinase expression (N=3) on LD day 6 upon MCUb silencing as compared to non targeting control. Densitometric analysis using ImageJ is presented below the blot. **(I)** qRT–PCR analysis showing decrease in MCUb mRNA expression upon MCUb silencing in LP primary human melanocytes (N=3). **(J)** Representative pellet pictures of siNT control and siMCUb in LP primary human melanocytes (N=3). **(K)** qRT–PCR analysis showing increase in GP100, Tyrosinase and DCT mRNA expression upon MCUb silencing in primary human melanocytes (N=3-4). **(L)** qRT–PCR analysis showing decrease in MICU1 mRNA expression upon MICU1 silencing in B16 cells (N=4). **(M)** Representative mitochondrial Ca^2+^ imaging traces of CEPIA2mt in siNT control and siMICU1 B16 cells stimulated with 100µM histamine. **(N)** Quantitation of resting mitochondrial Ca^2+^ with CEPIA2mt in siNT control and siMICU1 B16 cells where “n” denotes the number of ROIs. **(O)** Quantitation of mitochondrial Ca^2+^ uptake by calculating increase in CEPIA2mt signal (ΔCEPIA2mt) in siNT control and siMICU1 B16 cells upon stimulation with 100µM histamine where “n” denotes the number of ROIs. **(P)** Representative pellet pictures of siNT control and siMICU1 on LD day 6 (N=3). **(Q)** Melanin content estimation in siNT and siMICU1 B16 cells on LD day 6 (N=3). Data presented are mean ± S.E.M. For statistical analysis, unpaired student’s *t*-test was performed for panels C, D, N, O while one sample *t*-test was performed for panels A, F, I, K, L, Q using GraphPad Prism software. Here, * *p* <0.05; ** *p* < 0.01; *** *p* < 0.001 and **** *p* < 0.0001.

We next corroborated the role of MCUb in regulating melanogenesis in primary human melanocytes. To examine significance of MCUb in primary human melanocytes, we transiently silenced MCUb using siRNAs in lightly pigmented primary human melanocytes. We first validated MCUb knockdown in primary cells via qRT–PCR analysis. We observed around 60% decrease in the MCUb mRNA expression in the siMCUb as compared to siNT transfected primary human melanocytes (**Figure 3I**). The knockdown of MCUb resulted in the visible increase in melanogenesis as evident in pellet pictures of siMCUb in comparison to siNT control (**Figure 3J**). Further, we analyzed mRNA expression of key melanogenic enzymes and melanosomal structural proteins i.e. GP100, tyrosinase and DCT in the siNT and siMCUb cells. We observed that MCUb silencing results in the significant increase in the expression of GP100, tyrosinase and DCT (**Figure 3K**). These findings suggest that the increase in expression of key melanogenic enzymes and melanosome structural protein contribute to enhanced melanogenesis observed in MCUb silenced melanocytes. Taken together, the data from B16 LD model and primary melanocytes establish MCUb as a negative regulator of pigmentation.

To further strengthen the role of mitochondrial Ca^2+^ signaling in melanogenesis, we performed siRNA mediated silencing of another key MCU regulator MICU1 (Perocchi, Gohil et al., 2010) in B16 LD pigmentation model. MICU1 acts as a gatekeeper of the MCU as increase in Ca^2+^ levels is sensed by MICUs, which in turn stimulates opening of MCU channel pore (Mallilankaraman, Doonan et al., 2012). We characterized MICU1 siRNA using qRT–PCR analysis. We observed significant decrease in MICU1 mRNA levels in comparison to control siNT transfected B16 cells (**Figure 3L**). Next, we measured basal mitochondrial Ca^2+^ levels and histamine induced mitochondrial Ca^2+^ uptake upon MICU1 silencing. We observed a significant increase in resting mitochondrial Ca^2+^ levels (**Figure 3M and N**) as well as mitochondrial Ca^2+^ uptake upon MICU1 knockdown as compared to siNT control (**Figure 3M and O**). This data suggests that MICU1 silencing enhances mitochondrial Ca^2+^ signaling in our model system. Next, we evaluated pigmentation phenotype in B16 LD day6 cells. We observed a visible increase in LD day6 pigmentation as evident in the LD day6 pellet pictures upon knockdown of MICU1 as compared to siNT control (**Figure 3P**). We further quantitated the increase in pigmentation upon MICU1 silencing by performing melanin content assays. We observed that MICU1 silencing enhanced melanogenesis in B16 LD cells by ∼40% (**Figure 3Q**). Collectively, the data presented in **Figure 2** and **Figure 3** clearly demonstrate that mitochondrial Ca^2+^ positively regulates pigmentation.

### MCU regulates pigmentation *in vivo*

Next, we investigated the role of MCU in pigmentation *in vivo* by using zebrafish as a model system. Zebrafish are extensively used as a model system for pigmentation studies (Logan, Burn et al., 2006) as signaling events and molecular players involved in regulating pigmentation are largely conserved across vertebrates. Zebrafish embryos are transparent and develop rapidly. The pigmented melanophores, which are melanocyte equivalents in zebrafish appear within 36-48 hours post fertilization (hpf). In zebrafish, pigmentation analysis can be easily monitored visually and quantified with microscopic analysis as well as biochemical assays (Kelsh, Brand et al., 1996). Thus, zebrafish serves as an exciting model system to study pigmentation and examine the relevance of pigmentation regulators *in vivo*.

In first set of experiments, we employed morpholino (MO) based knockdown strategy. We used specific morpholinos (injected at single-cell stage) targeting zebrafish MCU and then followed changes in pigmentation. We first characterized the morpholinos targeting MCU by performing western blotting on zebrafish lysates at 48hrs post injection time point. We observed a significant decrease (around 60%) in the MCU protein expression with morpholino targeting zebrafish MCU as compared to control morpholino (**Figure 4A**). The quantitation of the extent of decrease in MCU protein levels upon MCU knockdown was performed on several independent experiments (**Supplementary Figure 3A**). Phenotypically, we observed a substantial reduction in the pigmented black-colored melanophores (melanocyte equivalents in zebrafish) in MCU morphants in comparison to the control embryos at 30hpf and 48hpf (hours post-fertilization) (**Figure 4B and C**). The decreased pigmentation phenotype was observed in 70% embryos at 30hpf (**Supplementary Figure 3B**). Next, we quantitated the decrease in melanogenesis upon MCU silencing by performing melanin content assays on zebrafish embryos. We found that the melanin content was decreased by ∼60% in MCU knockdown zebrafish embryos in comparison to control embryos (**Figure 4D**). To further corroborate *in vivo* data, we performed gain of function and rescue studies in zebrafish. We injected human MCU RNA in zebrafish for validating the role of MCU in pigmentation. We observed increase in the pigmented black-colored melanophores upon human MCU RNA injection as compared to the control embryos at 48 hpf (**Supplementary Figure 3C**). We quantitated the increase in pigmentation upon human MCU RNA injections in zebrafish by performing melanin content assays. Our quantitative analysis suggests that melanin content is increased by ∼40% upon human MCU RNA injections in comparison to control embryos (**Supplementary Figure 3D**). Moreover, we performed rescue experiments by injecting human MCU RNA along with MCU morpholinos. We observed rescue of hypopigmentation phenotype upon injection of human MCU RNA (**Figure 4E**) in MCU morphant embryos. The quantitative melanin content analysis on the zebrafish embryos clearly demonstrates that the decrease in pigmentation observed upon MCU knockdown in zebrafish can be rescued by co-injection of human MCU RNA (**Figure 4F**). Taken together, our zebrafish data elegantly establish an important role of MCU in regulating melanogenesis *in vivo*.

**Figure 4.**
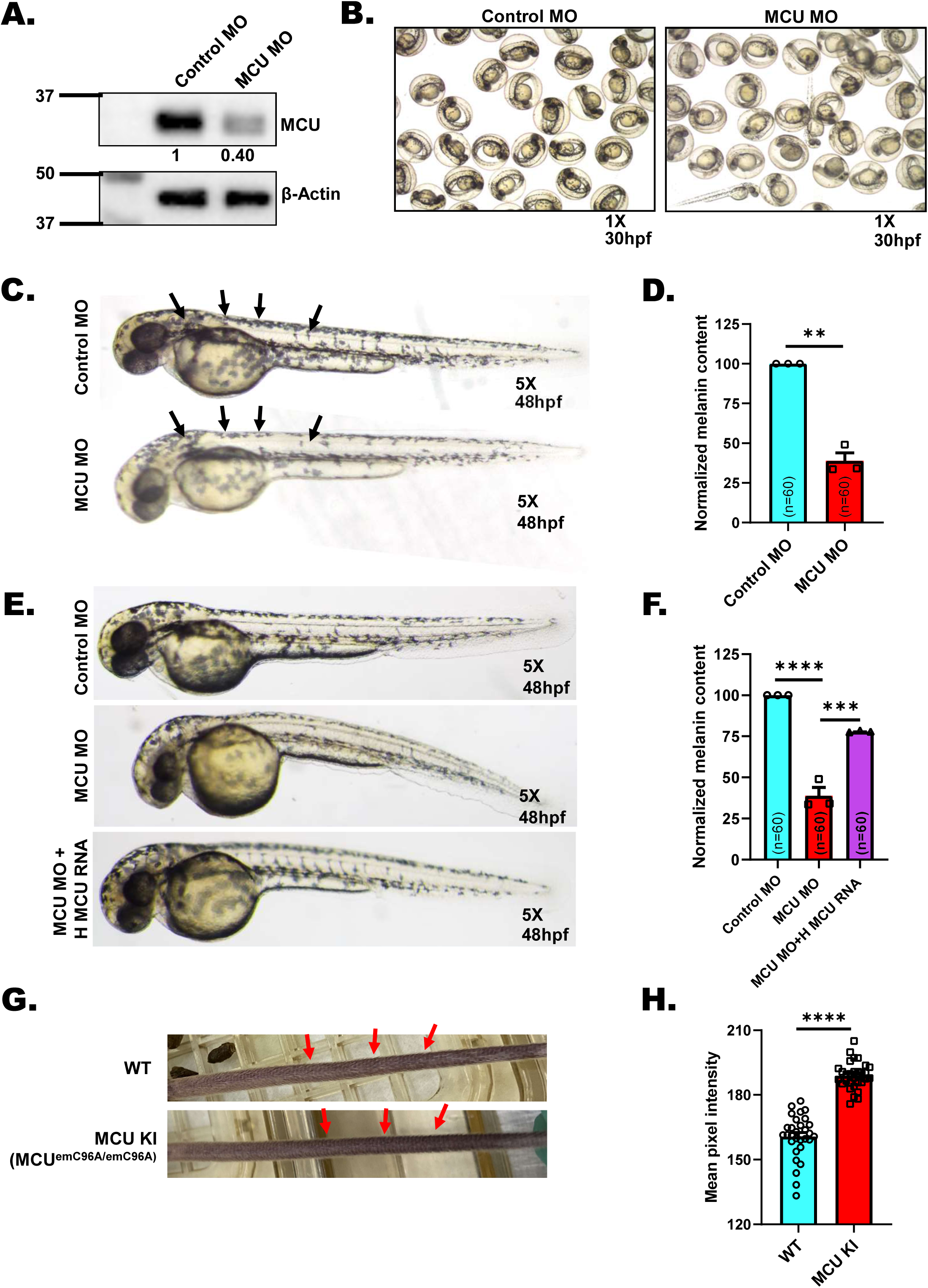
MCU regulates pigmentation *in vivo*. **(A)** Representative western blot showing expression of MCU in control MO and MCU MO. Densitometric analysis using ImageJ is presented below the blot (N=3). **(B)** Representative bright-field images of wild-type zebrafish embryos injected with either control morpholino or morpholino targeting zebrafish MCU at 30hpf (hours post fertilization) (N=3 independent experiments with ∼200 embryos/condition). **(C)** Representative bright-field images of wild-type zebrafish embryos injected with either control morpholino or morpholino targeting zebrafish MCU at 48hpf (hours post fertilization) (N=3 independent experiments with ∼200 embryos/condition). **(D)** Melanin-content estimation of control MO and MCU MO in zebrafish embryos in around 60 embryos from three independent sets of injections (N=3 independent experiments with 60 embryos/condition). **(E)** Representative bright-field images of zebrafish embryos injected with control morpholino; morpholino targeting zebrafish MCU and MCU morpholino injected with human MCU RNA (N=3 independent experiments with ∼200 embryos/condition). **(F)** Melanin-content estimation of control MO; MCU MO and MCU MO injected with human MCU RNA in zebrafish embryos in around 60 zebrafish embryos from three independent sets of injections (N=3 independent experiments with 60 embryos/condition). **(G)** Representative pictures of tail from 12 week-old wildtype and MCU-KI mice. **(H)** Quantitation of mean pixel intensity of tail pigmentation measured by ImageJ software in wild type mice and MCU KI mice (N=30 data points from 3 independent mice/condition) . Data presented are mean ± S.E.M. For statistical analysis, one sample *t*-test was performed for panel D; unpaired *t*-test was performed for panel H while one-way ANOVA followed by Tukey’s post-hoc test was performed for panel F using GraphPad Prism software. Here, ** *p* < 0.01; *** *p* < 0.001 and **** *p* < 0.0001.

Next, we assessed the role of MCU in mice epidermal pigmentation. For addressing this question, we utilized a recently reported MCU gain-of-function mouse model. This MCU C96A-knockin (KI) mice corresponds to human MCU C97A and results in hyperactivation of MCU channel activity (Tomar, Jana et al., 2019). It was demonstrated that MCU-mediated mitochondrial Ca^2+^ uptake rate was higher in this MCU-C96A KI mice (Tomar et al., 2019). Excitingly, we observed enhanced epidermal pigmentation in tail of these MCU-KI mice in comparison to wild type mice as pointed in the mice tail pictures presented in **Figure 4G**. It is important to highlight that epidermal pigmentation in mice is typically studied in mice tail as mice fur pigmentation is equivalent to human hair pigmentation. We further quantitated the difference in epidermal pigmentation between wild type and MCU KI mice with a standard protocol used for calculating pigmentation in animal models (Sultan et al., 2022). Briefly, we calculated mean pixel intensity of the tail pictures using ImageJ software. Notably, we observed a significant increase in the epidermal tail pigmentation in MCU-KI mice in comparison to wild type mice (**Figure 4H**). This data further validates that MCU plays a critical role in pigmentation *in vivo*. Collectively, our data from four independent model systems i.e. 1) B16 cells; 2) primary human melanocytes; 3) Zebrafish and 4) Mouse model clearly establish that mitochondrial Ca^2+^ signaling plays a crucial role in regulating vertebrate pigmentation.

### Unbiased transcriptome analysis identifies keratin filaments as key melanogenic regulators working downstream of mitochondrial Ca^2+^signaling

To identify the molecular mechanism regulating pigmentation downstream of mitochondrial Ca^2+^, we performed transcriptome profiling (RNA sequencing). RNA seq was performed on siNT, siMCU and siMCUb LD day6 cells, (**Figure 5A**) using Illumina NovaSeq. The data quality was checked using FastQC and MultiQC software. The QC passed reads were mapped onto indexed Mus musculus genome (GRCm39) using STAR v2 aligner. Gene level expression values were obtained as read counts using feature-counts software. Next, we carried out differential expression analysis using the DESEq2 package and normalized counts were obtained. The fold change was calculated for siMCU and siMCUb samples with respect to siNT control. Genes showing expression more than log2 fold change +1 with respect to siNT control were taken as upregulated genes in siMCU and siMCUb. While a cut off of log2 fold change less than -1 was set for determining downregulated genes in siMCU and siMCUb. First of all, we examined the levels of MCU and MCUb in the RNA seq data. As expected, we observed a significant decrease in their levels in the respective siRNA condition (**Figure 5B**). We then validated knockdown of MCU and MCUb in RNA seq samples by performing qRT-PCR analysis (**Figure 5C**). As a positive control, we checked expression of pigmentation genes upon knockdown of MCU and MCUb in RNA seq samples. We observed increase in expression of pigmentation genes upon knockdown of MCUb while MCU silencing led to decrease in expression of some of the key pigmentation genes (**Figure 5D**). Taken together, this initial scrutiny gave us confidence that our RNA sequencing is performed and analyzed appropriately.

**Figure 5.**
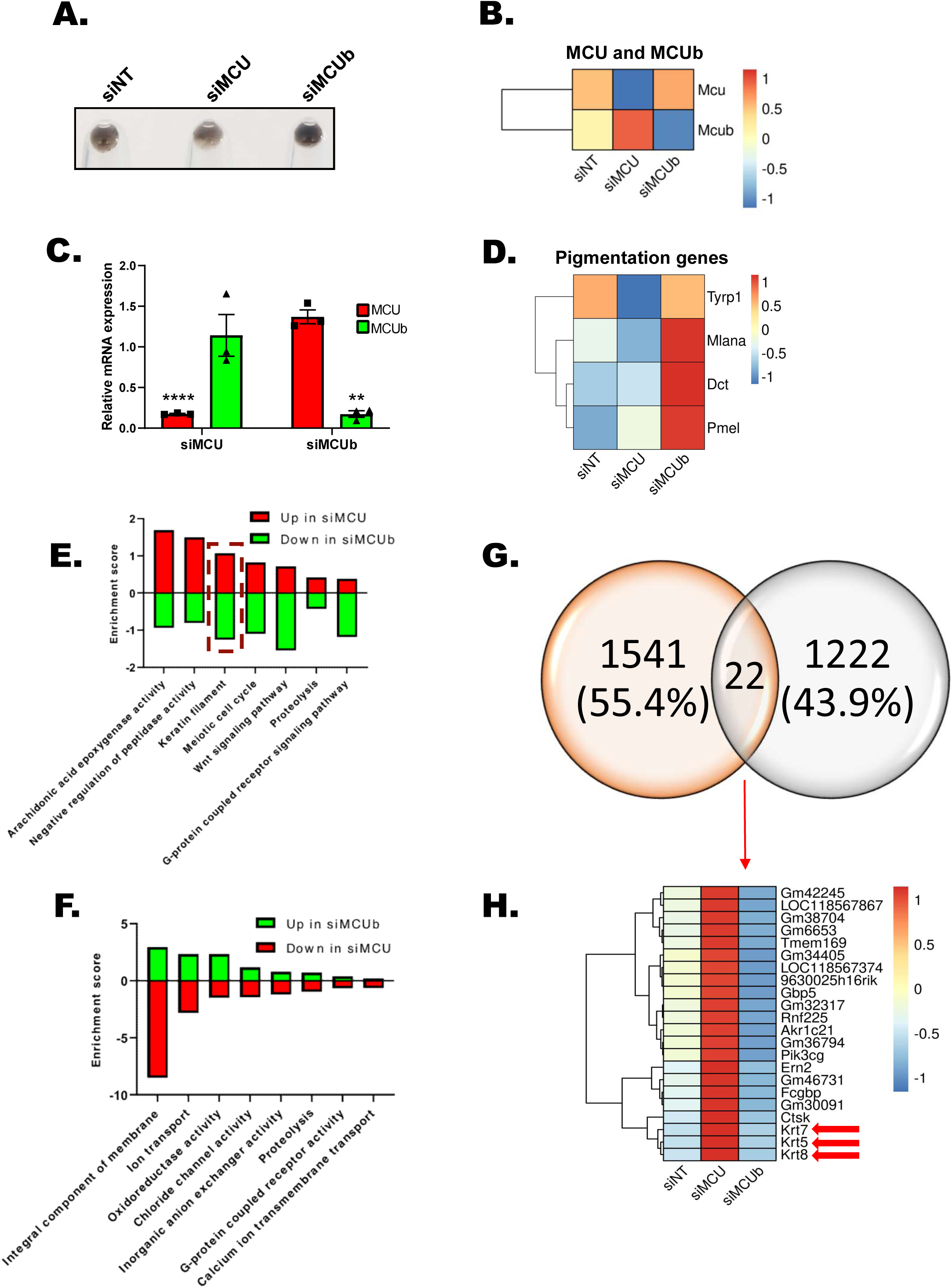
Transcriptomics for identifying the molecular mechanism connecting mitochondrial Ca^2+^ dynamics to melanogenesis. **(A)** Representative pellet pictures of B16 LD cells used for RNA sequencing. **(B)** Heatmap representing the expression of MCU and MCUb upon silencing of MCU and MCUb respectively. Scale from blue to red represents z-score for fold change from -1 to +1. **(C)** qRT–PCR analysis showing decrease in MCU and MCUb mRNA expression upon MCU and MCUb silencing respectively (N=3). **(D)** Heatmap representing the expression of pigmentation genes upon silencing of MCU and MCUb respectively. Scale from blue to red represents z-score for fold change from -1 to +1. **(E)** Common oppositely regulated pathways up in siMCU and down in siMCUb **(F)** Common oppositely regulated pathways down in siMCU and up in siMCUb. **(G)** Venn Diagram representing common differentially regulated genes upon silencing of MCU and MCUb **(H)** Heatmap representing the expression of 22 common differentially regulated genes upon silencing of MCU and MCUb respectively. Scale from blue to red represents z-score for fold change from -1 to +1. Data presented are mean ± S.E.M. For statistical analysis, one sample *t*-test was performed for panel C using GraphPad Prism software. Here, ** *p* < 0.01and **** *p* < 0.0001.

In order to identify the pathways that are modulated upon MCU and MCUb silencing during pigmentation, we performed pathway enrichment analysis on upregulated and downregulated genes (in siMCU and siMCUb conditions), using DAVID software. Pathway enrichment analysis was performed based on the Gene Ontology (GO) database. Enriched pathways were plotted with enrichment scores in Graphpad. Since we observed opposing pigmentation phenotypes upon MCU and MCUb silencing, we adopted the strategy of determining the pathways that are oppositely regulated in siMCU and siMCUb conditions. We plotted the common but oppositely modulated pathways i.e. up in siMCU and down in siMCUb and vice versa (down in siMCU and up in siMCUb) (**Figure 5E and F**). Top oppositely regulated pathways up in siMCU and down in siMCUb were arachidonic acid epoxygenase activity, keratin filament, Wnt signaling, G-protein coupled receptor signaling pathway and meiotic cell cycle among others. Likewise, top common but oppositely regulated pathways down in siMCU and up in siMCUb were integral component of membrane, ion transport, oxidoreductase activity and chloride channel activity.

Next, to identify common but oppositely regulated genes, we plotted Venn diagram for genes up in siMCU and down in siMCUb. We observed 22 such differentially regulated genes upon silencing of MCU and MCUb (**Figure 5G**). Interestingly 3 out of 22 differentially regulated genes belong to keratin intermediate filament family i.e. keratin 5, 7 and 8 (**Figure 5H**). Upon detailed literature survey we found that the role of keratins in regulating melanogenesis remains poorly understood. Two independent groups have shown association of keratins with pigmentation in sheep and alpaca skins (Cui, Song et al., 2016; Fan, Xie et al., 2013). Keratin and keratin related proteins were highly expressed in white sheep skin as compared to black sheep skin (Fan et al., 2013). In another study expression of keratin 2 was higher in brown alpaca skin in comparison to white alpaca skin (Cui et al., 2016). Additionally, the authors reported that ectopic overexpression of keratin 2 in sheep melanocytes results in increase in melanogenesis thereby suggesting that keratin 2 regulates melanogenesis in sheep. Further, literature review for role of keratins in human skin diseases highlighted an association of keratin 5 mutations with hyper-pigmentary skin disorders such as Dowling Degos disease and epidermolysis bullosa simplex with mottled pigmentation (Betz, Planko et al., 2006; Moog, de Die-Smulders et al., 1999). Another recent study demonstrated that the expression of keratin 5 is higher in hyper pigmentary conditions such as senile lentigo, photo exposed skin and Dowling Degos disease in comparison to normal mammary skin (Cario, Pain et al., 2020). The authors further showed that keratin 5 levels are lower in case of depigmented vitiligo skin (Cario et al., 2020). These clinical studies establish an important association between keratin 5 mutations/expression profile with pigmentary disorders. However, there is no study that clearly explains the functional significance of keratin 5 in pigmentary disorders. Further, how is keratin 5 expression regulated in these pigmentary disorders remains unappreciated. Therefore, based on pathway enrichment analysis and literature survey we focused on understanding the role of keratin filaments in regulating melanogenesis downstream of mitochondrial Ca^2+^ signaling.

### Keratin 5 regulates melanogenesis and mitochondrial Ca^2+^ uptake

To understand the role of keratin 5, 7 and 8 in pigmentation, we started by characterizing the siRNAs targeting them. As presented in **Supplementary Figure 4A-C**, siRNAs against keratin 5, 7 and 8 were able to decrease the target mRNAs by ∼75% in B16 cells. Next, we used these siRNAs to decrease the keratins expression in B16 cells during LD pigmentation. We performed siRNA mediated silencing on LD day3 and terminated the experiment on LD day6. The knockdown of all three keratins led to a substantial phenotypic decrease in LD pigmentation as evident in the LD day6 pellet pictures (**Figure 6A, C and E**). We further quantitated the decrease in pigmentation by performing melanin content assays. We observed that silencing of keratin 5, 7 and 8 decreased melanogenesis by over 50% in B16 cells (**Figure 6B, D and F**). Hence, our data suggests that keratin 5, 7 and 8 positively regulate B16 LD melanogenesis. Since all three keratins (keratin 5, 7 and 8) gave similar pigmentation phenotype and clinically keratin 5 is associated with pigmentary disorders, we focused on keratin 5 for subsequent studies. We first validated role of keratin 5 in regulating pigmentation in primary human melanocytes. We silenced keratin 5 using human keratin 5 siRNA in primary human melanocytes. We examined keratin 5 knockdown by performing qRT–PCR analysis and observed a drastic decrease (around 80%) in keratin 5 mRNA levels in sikeratin 5 condition in comparison to control siNT (**Figure 6G**). The knockdown of keratin 5 resulted in visible decrease in melanogenesis as evident in the pellet pictures (**Figure 6H**). These observations were rather surprising as the expression of keratin 5 increases upon MCU silencing and decreases upon MCUb knockdown (**Figure 5E and H**) and therefore we were initially expecting that silencing of keratin 5 would result in increase in melanogenesis as observed in case of siMCUb. This data suggests that keratin 5 could be playing some sort of compensatory/feedback function downstream of mitochondrial Ca^2+^ signaling. Recent literature suggests that keratin filaments can modulate mitochondrial organization and function (Schwarz & Leube, 2016; Silvander, Kvarnstrom et al., 2017; Steen, Chen et al., 2020). However, role of keratins in regulating mitochondrial Ca^2+^ dynamics remains unappreciated. Therefore, we investigated role of keratin 5 in controlling mitochondrial Ca^2+^ uptake. We first cloned human keratin 5 in a mcherry vector and confirmed that transfection of mcherry tagged keratin 5 leads to a robust increase in the expression of keratin 5 in B16 cells at mRNA levels (**Figure 6I**). Next, we measured resting mitochondrial Ca^2+^ levels and histamine induced mitochondrial Ca^2+^ uptake in empty vector (EV) control and keratin 5 overexpression conditions. Interestingly, we observed that although basal mitochondrial Ca^2+^ levels are comparable in two conditions, mitochondrial Ca^2+^ uptake was significantly increased upon keratin 5 overexpression (**Figure 6J-L**). This suggests that the increase in keratin 5 expression observed upon MCU knockdown could be a compensatory mechanism of the cells to counteract the adverse effects associated with dysregulation of mitochondrial Ca^2+^ homeostasis. We further corroborated our results by examining basal mitochondrial Ca^2+^ levels and mitochondrial Ca^2+^ uptake upon keratin 5 silencing. We found that keratin 5 knockdown in B16 cells results in a substantial decrease in mitochondrial Ca^2+^ uptake but there is no significant change in the resting mitochondrial Ca^2+^ levels (**Figure 6M-O**). Taken together, our data demonstrates that MCU complex driven mitochondrial Ca^2+^ signaling and keratin 5 function in a classical feedback loop wherein mitochondrial Ca^2+^ dynamics regulates keratin 5 expression while keratin 5 in turn modulates MCU mediated mitochondrial Ca^2+^ uptake (**Figure 6P**).

**Figure 6.**
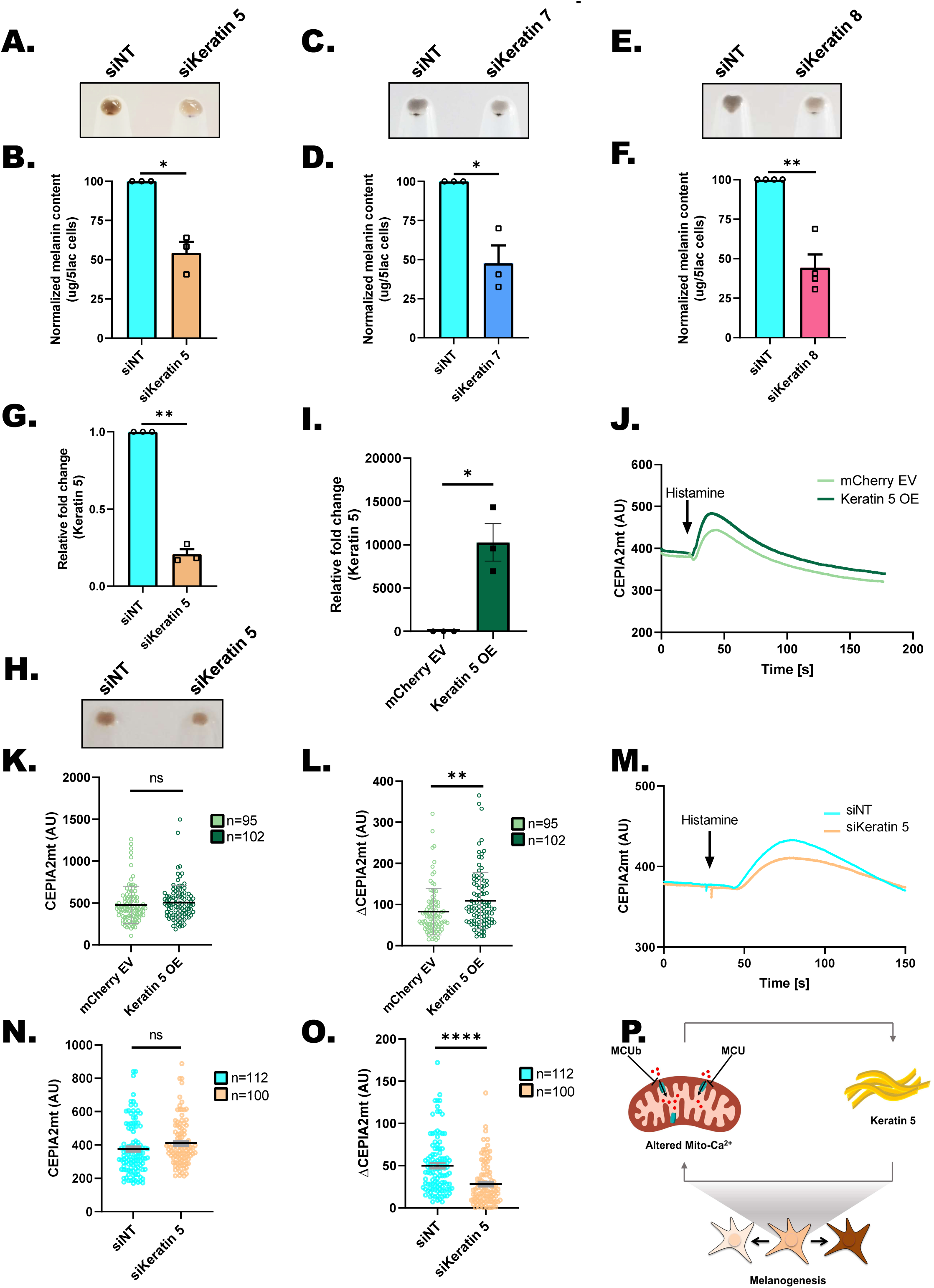
Keratins regulate melanogenesis. **(A)** Representative pellet pictures of siNT control and siKeratin 5 on LD day 6 (N=3). **(B)** Melanin content estimation of siNT and siKeratin 5 B16 cells on LD day 6 (N=3). **(C)** Representative pellet pictures of siNT control and siKeratin 7 on LD day 6 (N=3). **(D)** Melanin content estimation of siNT and siKeratin 7 B16 cells on LD day 6 (N=3). **(E)** Representative pellet pictures of siNT control and siKeratin 8 on LD day 6 (N=4). **(F)** Melanin content estimation of siNT and siKeratin 8 B16 cells on LD day 6 (N=4). **(G)** qRT–PCR analysis showing decrease in Keratin 5 mRNA expression upon Keratin 5 silencing in LP primary human melanocytes (N=3). **(H)** Representative pellet pictures of siNT control and siKeratin 5 in LP primary human melanocytes (N=3). **(I)** qRT–PCR analysis showing increase in Keratin 5 mRNA expression upon ectopic expression of Keratin5 in B16 cells (N=3). **(J)** Representative mitochondrial Ca^2+^ imaging traces of CEPIA2mt in mCherry empty vector (EV) control and Keratin 5 overexpressing (OE) cells stimulated with 100µM histamine. **(K)** Quantitation of resting mitochondrial Ca^2+^ with CEPIA2mt in mCherry empty vector (EV) control and Keratin 5 overexpressing (OE) cells where “n” denotes the number of ROIs. **(L)** Quantitation of mitochondrial Ca^2+^ uptake by calculating increase in CEPIA2mt signal (ΔCEPIA2mt) in mCherry empty vector (EV) control and Keratin 5 overexpressing (OE) cells upon stimulation with 100µM histamine where “n” denotes the number of ROIs. **(M)** Representative mitochondrial Ca^2+^ imaging traces of CEPIA2mt in siNT control and siKeratin 5 cells stimulated with 100µM histamine. **(N)** Quantitation of resting mitochondrial Ca^2+^ with CEPIA2mt in siNT control and siKeratin 5 cells where “n” denotes the number of ROIs. **(O)** Quantitation of mitochondrial Ca^2+^ uptake by calculating increase in CEPIA2mt signal (ΔCEPIA2mt) in siNT control and siKeratin 5 cells upon stimulation with 100µM histamine where “n” denotes the number of ROIs. **(P)** Mitochondrial Ca^2+^ signaling and keratin 5 regulate each other in a feedback loop to ensure optimal melanogenesis. Data presented are mean ± S.E.M. For statistical analysis, one sample *t*-test was performed for panels B, D, F, G, I while unpaired *t*-test was performed for panel K, L, N, O using GraphPad Prism software. Here, * *p* <0.05; ** *p* < 0.01; *** *p* < 0.001 and **** *p* < 0.0001.

### Keratin 5 regulates melanosome maturation and positioning

We next investigated if keratin 5 works downstream of mitochondrial Ca^2+^ signaling to regulate melanogenesis. We therefore examined if keratin 5 can modulate MCUb knockdown induced hyperpigmentation phenotype. We performed either MCUb silencing alone or along with keratin 5 knockdown in the LD pigmentation model. As expected, MCUb silencing enhanced pigmentation and keratin 5 knockdown in MCUb silenced cells led to rescue of the pigmentation phenotype and corresponding melanin content (**Figure 7A and B**). To understand the molecular mechanism driving the rescue of pigmentation phenotype, we studied expression and activity of key melanogenic proteins upon MCUb plus keratin 5 knockdown and compared it with MCUb silencing alone. As presented in **Figure 7C**, co-silencing of keratin 5 plus MCUb inhibits the increase in GP100 and DCT expression observed in siMCUb condition.

**Figure 7.**
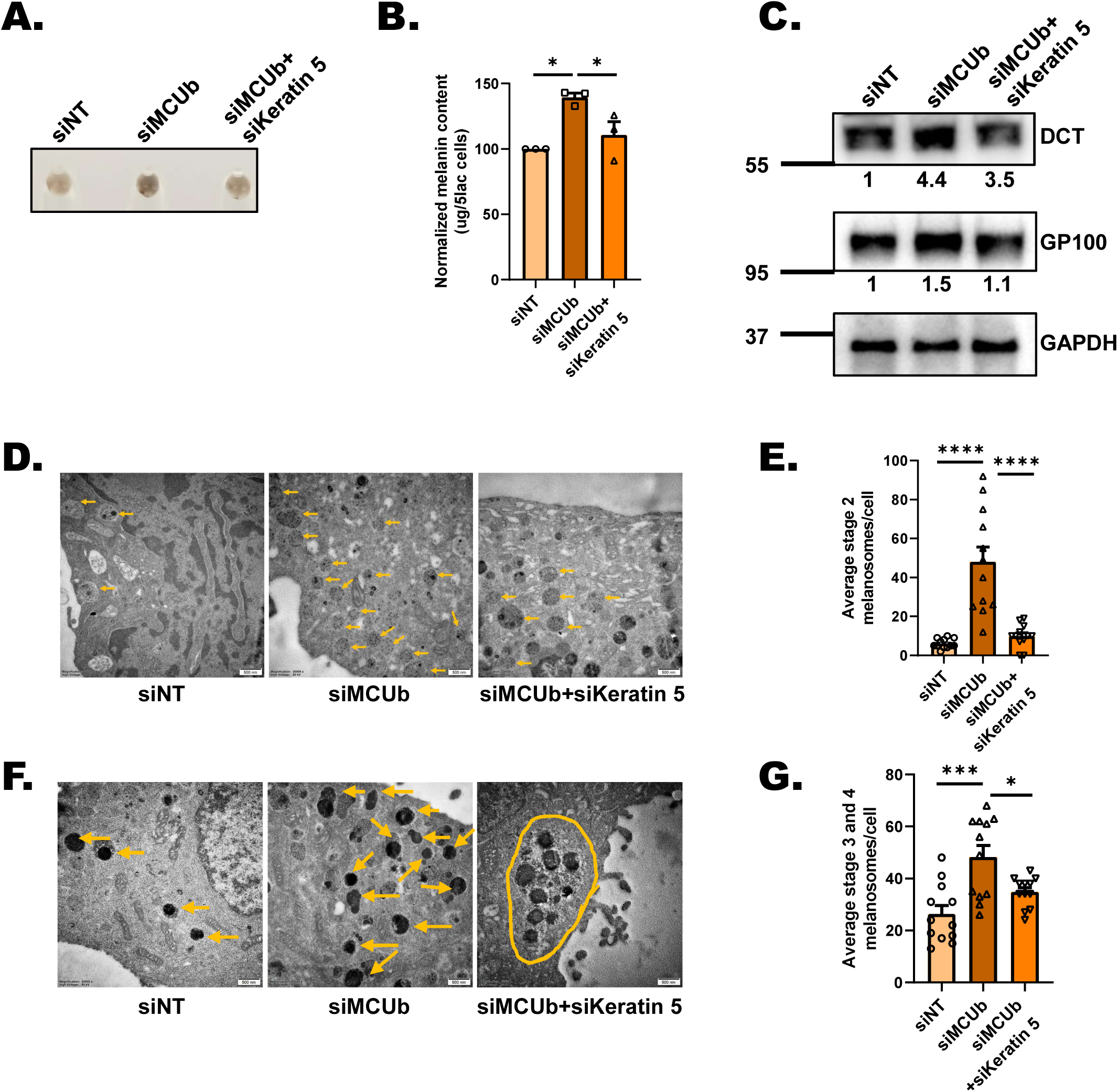
Keratin 5 regulates melanosome maturation and positioning. **(A)** Representative pellet pictures of siNT control, siMCUb and siMCUb+ siKeratin 5 on LD day 6 (N=3). **(B)** Melanin content estimation of siNT, siMCUb and siMCUb+ siKeratin 5 on LD day 6 (N=3). **(C)** Representative western blot showing expression of DCT and GP100 on LD day 6 upon MCUb, siMCUb+ siKeratin 5 silencing as compared to non-targeting control. Densitometric analysis using ImageJ is presented below the blot. **(D)** TEM images of siNT control, siMCUb and siMCUb+ siKeratin 5 B16 LD day6 cells. Yellow arrows correspond to stage 2 melanosomes in these cells on LD day6. **(E)** Quantification of number of stage 2 melanosomes/cell in siNT control, siMCUb and siMCUb+ siKeratin 5 B16 LD day6 cells (N= 12 cells/condition). **(F)** TEM images of siNT control, siMCUb and siMCUb+ siKeratin 5 B16 LD day6 cells. Yellow arrows correspond to melanin rich mature (stage 3 and 4) melanosomes in these cells on LD day6. Yellow oval shape shows clustering of melanosomes in siMCUb+ siKeratin 5. **(G)** Quantification of number of mature melanosomes/cell in siNT control, siMCUb and siMCUb+ siKeratin 5 B16 LD day6 (N= 12 cells/condition). Data presented are mean ± S.E.M. For statistical analysis, one-way ANOVA followed by Tukey’s post-hoc test was performed using GraphPad Prism software. Here, * *p* <0.05; *** *p* < 0.001 and **** *p* < 0.0001.

Finally, we carried out ultrastructural studies using transmission electron microscope (TEM) for investigating the effect of MCUb knockdown and MCUb plus keratin 5 silencing on melanosome biogenesis, maturation and positioning. We observed that MCUb silencing led to a significant increase in melanosome biogenesis (i.e. number of II melanosomes) (**Figure 7D and E**) and maturation (i.e. number of stage III and IV melanosomes) (**Figure 7F and G**). While simultaneous knockdown of MCUb and keratin 5 leads to a substantial decrease in melanosome biogenesis (**Figure 7D and E**) and maturation as compared to MCUb silencing alone (**Figure 7F and G**). Further, in case of MCUb silencing alone melanosomes were distributed throughout cells whereas upon MCUb plus keratin 5 knockdown melanosomes were typically clustered together in clumps (**Figure 7F**). It is important to highlight that similar clustering of melanosomes is reported in case of human samples of epidermolysis bullosa simplex with mottled pigmentation (Moog et al., 1999), a clinical condition associated with keratin 5 mutations. Taken together, our data suggests that mitochondrial Ca^2+^ signaling regulates keratin 5 expression in melanocytes and that in turn modulates melanosome biogenesis, maturation and positioning.

### MCU drives Keratin 5 expression via NFAT transcription factor

We next explored the possible molecular choreography that connects keratin 5 expression to mitochondrial Ca^2+^ dynamics. Since we observed differential mRNA expression of keratin 5 upon silencing of MCU and MCUb, we surveyed the literature regarding transcription factors directly activated upon changes in mitochondrial Ca^2+^ signaling. Interestingly, we came across a recent study that demonstrated NFAT (Nuclear Factor of Activated T cells) transcription factors’ activation downstream of mitochondrial Ca^2+^ dynamics (Yoast, Emrich et al., 2021). The authors elegantly demonstrated that MCU silencing leads to NFAT1/4 activation and their nuclear translocation (Yoast et al., 2021). Based on this study, we bioinformatically scanned the keratin 5 promoter for potential NFAT binding sites. We used multiple bioinformatics tools, namely ContraV3 (Kreft, Soete et al., 2017), EPD-Search Motif Tool (Dreos, Ambrosini et al., 2015), PSCAN (Zambelli, Pesole et al., 2009) and JASPAR 2022 (Castro-Mondragon, Riudavets-Puig et al., 2022), to scan keratin 5 promoter for putative binding sites for Ca^2+^ activated NFAT1-4 transcription factors.

We observed that there are no conversed sites for NFAT3 in keratin 5 promoter. Although there are couple of potential binding sites for NFAT1 and NFAT4 in keratin 5 promoter, they are neither predicted by all four bioinformatics tools with confidence nor they are highly conserved. While our survey for NFAT2 binding site (**Figure 8A**) on keratin 5 promoter led to identification of 4 highly conserved sites (**Figure 8B**). These putative sites were predicted by all four bioinformatics tools with high confidence. This suggests that MCU can potentially control transcriptional regulation of keratin 5 through increased activation of NFAT2. Our RNA sequencing data showed that along with keratin 5, keratin 7 and 8 are also differentially regulated upon changes in mitochondrial Ca^2+^ dynamics. Therefore, we also examined keratin 7 and 8 promoters for putative NFAT2 binding sites. Indeed, we found presence of 6 and 2 conserved as well as high confidence NFAT2 binding sites in keratin 7 and 8 promoters respectively (**Supplementary Figure 5A-D**).

**Figure 8.**
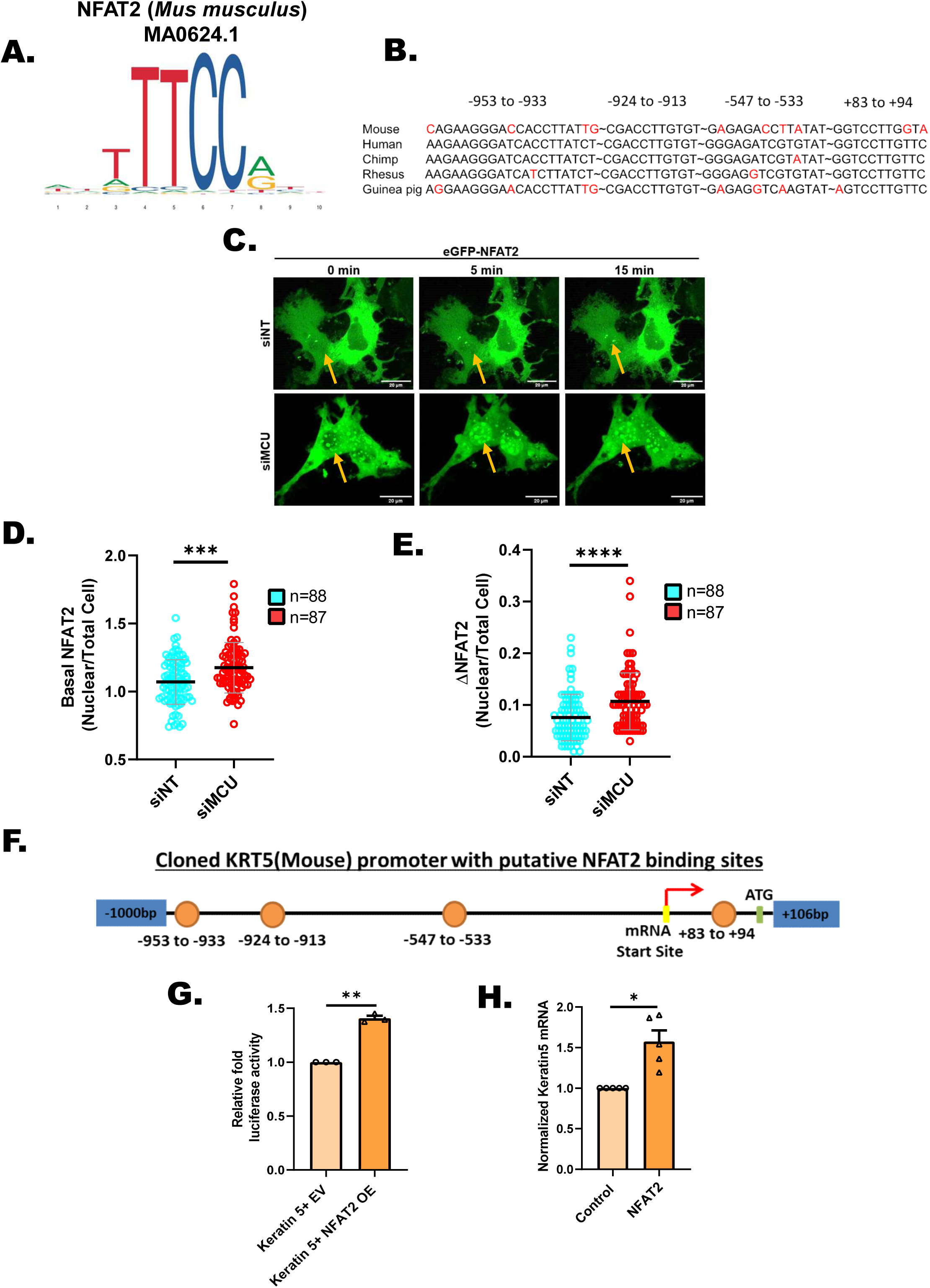
NFAT2 connects MCU to Keratin 5 expression. **(A)** Position weight matrix of mouse NFAT2 consensus binding sequence. **(B)** Multispecies sequence alignment of putative NFAT2 binding sites in the mouse keratin 5 (KRT5) core promoter. **(C)** Representative confocal microscopy images of differential NFAT2 nuclear translocation in B16 cells transfected with siNT or siMCU, upon stimulation with 100uM histamine. Images have been captured at 63X(oil) magnification in the GFP channel in a Zeiss confocal microscope. Images presented are maximum intensity projections. **(D)** Quantification of basal NFAT2 nuclear levels in B16 cells with MCU knockdown. Live cell imaging was performed in B16 cells transfected with siMCU or control siRNA along with eGFP-NFAT2 overexpression. Nuclear/Cytoplasmic ratio of NFAT2 was quantified by determining the GFP fluorescence from the nucleus versus total area of a single cell. ‘n’ represents the total number of cells imaged per condition. **(E)** Quantification of NFAT2 nuclear translocation upon histamine stimulation. Live cell imaging was performed in B16 cells transfected with siMCU or control siNT along with eGFP-NFAT2 overexpression. Nuclear/Cytoplasmic ratio of NFAT2 was quantified by determining the GFP fluorescence from the nucleus versus total area of a single cell. NFAT2 nuclear translocation in response to histamine (100µM) stimulation was quantified (5min and 15min post stimulation) by calculating the difference between maximal NFAT2 nuclear/total ratio upon histamine stimulation and basal NFAT2 nuclear/total ratio of unstimulated cells. ‘n’ represents the total number of cells imaged per condition. **(F)** Schematic representation of putative NFAT2 binding sites in the mouse KRT5 core promoter cloned in pGL4.23 luciferase vector. **(G)** Normalized luciferase activity of the KRT5 promoter luciferase reporter vector with overexpression of eGFP-NFAT2 or empty vector control in B16 cells (N=3). **(H)** Normalized mRNA expression of mouse KRT5 gene in B16 cells with overexpression of eGFP-NFAT2 or empty vector control (N=6). Data presented are mean ± S.E.M. For statistical analysis, one sample *t*-test was performed for panels G, H while Mann-Whitney Test was performed for panels D, E using GraphPad Prism software. Here, * *p* <0.05; ** *p* < 0.01; *** *p* < 0.001 and **** *p* < 0.0001.

To study the role of NFAT2 in connecting mitochondrial Ca^2+^ signaling to keratin 5 expression, first of all we investigated NFAT2 activation upon MCU silencing in B16 cells. As nuclear translocation is a prerequisite of NFAT transcriptional activity, we examined the effects of MCU knockdown on NFAT2 nuclear translocation. We overexpressed GFP tagged NFAT2 in B16 cells with MCU knockdown and performed live cell imaging to study NFAT2 nuclear translocation (**Figure 8C**). We observed an increase in NFAT2 nuclear translocation in MCU knockdown cells as compared to control siRNA condition under resting/basal conditions (**Figure 8C and D**). This implies that knockdown of MCU reconfigures the cellular Ca^2+^ homeostasis leading to NFAT2 activation and nuclear translocation. Further, we found that upon stimulation with histamine, there was a significant enhancement in NFAT2 nuclear translocation in MCU knockdown cells in comparison to control cells (**Figure 8C and E**). This clearly demonstrates that reduction in MCU expression results in increased NFAT2 nuclear translocation, which could in turn enhance keratin 5 gene transcription.

To evaluate the functional significance of NFAT2 in regulating keratin 5 transcription and promoter activity, we cloned keratin 5 promoter with putative NFAT2 binding sites in a luciferase vector (**Figure 8F**). We performed *in vitro* luciferase assay with this keratin 5 promoter along with NFAT2 overexpression in B16 cells. Indeed, we observed a significant increase in luciferase activity of the keratin 5 promoter upon NFAT2 overexpression (**Figure 8G**). This clearly indicated that NFAT2 acts as a transcriptional regulator of keratin 5 expression. Further, we examined mRNA expression of keratin 5 upon NFAT2 overexpression. As expected, we observed an evident increase in keratin 5 mRNA expression validating a positive regulation of keratin 5 gene expression by NFAT2 (**Figure 8H**). Taken together, our extensive bioinformatics analysis, keratin 5 promoter reporter assays and mRNA expression data clearly demonstrates that NFAT2 acts as a bridging transcription factor that connects MCU silencing to keratin 5 gene expression. It is important to note that MCU silencing concomitantly decreases pigmentation and enhances transcription of keratin 5, a positive regulator of melanogenesis. A possible explanation for this observation could be that increase in keratin 5 levels act as a negative feedback loop for ensuring that melanogenesis is not severely compromised upon perturbations in mitochondrial Ca^2+^ signaling.

### Mitoxantrone inhibits physiological pigmentation

Next, we investigated the potential of targeting MCU mediated mitochondrial Ca^2+^ uptake to modulate pigmentation. We used mitoxantrone, an FDA approved drug which is highly specific MCU blocker (Arduino, Wettmarshausen et al., 2017), to examine the significance of blocking MCU in managing melanogenesis. We used physiological melanogenic stimuli αMSH for inducing melanogenesis in B16 cells and co-treated cells with mitoxantrone to evaluate contribution of MCU in αMSH stimulated pigmentation. We utilized αMSH for carrying out these studies as αMSH induces ER Ca^2+^ release and enhances cytosolic Ca^2+^ levels (Motiani et al., 2018). This in turn would lead to mitochondrial Ca^2+^ influx and mitoxantrone will block MCU mediated mitochondrial Ca^2+^ uptake. Excitingly, we observed that mitoxantrone treatment results in substantial reduction in αMSH stimulated physiological melanogenesis (**Figure 9A**). We further quantitated these phenotypic changes by measuring mean pixel intensity of pellets using ImageJ software and observed that the mitoxantrone treatment leads to around 70% decrease in αMSH induced melanogenesis (**Figure 9B**). This data suggests that MCU mediated mitochondrial Ca^2+^ influx can be targeted for modulating pigmentation.

**Figure 9.**
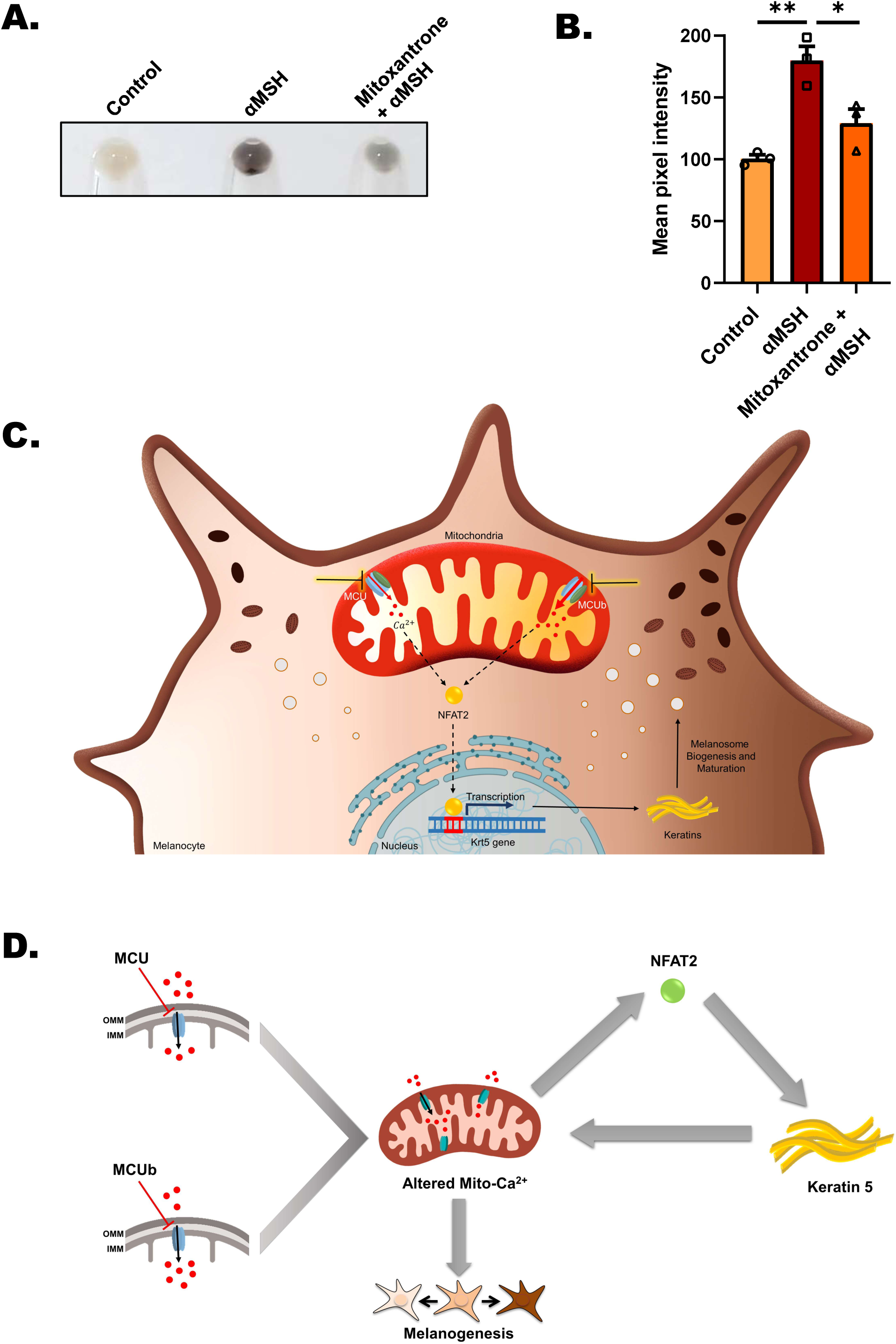
Mitoxantrone inhibits *α*MSH-induced physiological pigmentation. **(A)** Representative pellet pictures upon αMSH, αMSH+mitoxantrone treatment in B16 cells as compared to vehicle control. **(B)** Mean pixel intensity of αMSH, αMSH+mitoxantrone treatment in B16 cells as compared to vehicle control (N=3). **(C)** Graphical summary of the work illustrating that silencing of MCU decreases melanogenesis while MCUb knockdown enhances melanogenesis. Transcription factor NFAT2 gets activated upon MCU silencing and that in turn induces keratin 5 expression. Keratin 5 drives melanogenesis by augmenting melanosome biogenesis and maturation. **(D)** Diagrammatic representation of negative feedback loop operating via MCU-NFAT2 Keratin 5 signaling module to maintain mitochondrial Ca^2+^ homeostasis and melanogenesis. Data presented are mean ± S.E.M. For statistical analysis, one-way ANOVA followed by Tukey’s post-hoc test was performed for panel B using GraphPad Prism software. * *p* <0.05; ** *p* < 0.01.

In summary, we have identified a *de novo* critical role of mitochondrial Ca^2+^ dynamics in driving pigmentation using several independent *in vitro* and *in vivo* model systems. Our data demonstrates that mitochondrial Ca^2+^ signaling is a crucial regulator of melanosome biogenesis and maturation. Further, by performing unbiased RNA seq analysis, we have deciphered the molecular mechanism that connects mitochondrial Ca^2+^ signaling to melanogenesis. We report that mitochondrial Ca^2+^ signaling regulates transcription of keratin filaments via NFAT2 transcription factor. The keratin filaments in turn contribute to melanosome biogenesis and maturation (**Figure 9C**). Importantly, we report that keratin 5 can regulate mitochondrial Ca^2+^ uptake. Therefore, MCU-NFAT2-Keratin 5 signaling cascade works as a negative feedback loop that maintains mitochondrial Ca^2+^ homeostasis and ensures optimal pigmentation (**Figure 9D**). Finally, we demonstrate that blocking of MCU results in inhibition of pigmentation. Thus highlighting the potential of targeting MCU for clinical management of pigmentary disorders.

## Discussion

Ionic homeostasis including intracellular Ca^2+^ dynamics plays an important role in driving human skin pigmentation (Bellono & Oancea, 2014; Sharma, Sharma et al., 2023). Physiologically relevant doses of UV light induces rise in cytosolic Ca^2+^ levels, which leads to enhanced melanin synthesis (Bellono, Kammel et al., 2013). Similarly, rise in free cytosolic Ca^2+^ upon VDAC1 silencing augments melanogenesis by activating MITF mediated transcription of melanogenic genes (Wang, Gong et al., 2022). These studies demonstrate that increase in cytosolic Ca^2+^ levels promotes melanogenesis. Recently, mitochondria localized K^+^-dependent Na^+^/Ca^2+^ exchanger 5 (NCKX5), also known as SLC24A5, was reported to regulate melanosome biogenesis and melanin synthesis (Zhang et al., 2019). It was demonstrated that NCKX5 controls melanosome Ca^2+^ levels and that in turn regulates melanosome maturation and melanogenesis (Zhang et al., 2019). Further, our earlier work demonstrated that physiological melanogenic stimuli αMSH increases cytosolic Ca^2+^ levels by stimulating ER Ca^2+^ release (Motiani et al., 2018). This in turn activates STIM1 oligomerization and enhances MITF driven melanogenesis. Interestingly, we recently reported that MITF transcriptionally regulates STIM1 expression in melanocytes and their expression concomitantly increases in sun-exposed tanned human skin (Tanwar et al., 2022b). This suggests that a positive feed forward loop driven by ER Ca^2+^ release is a critical regulator of melanogenesis. In cellular systems, a major proportion of ER Ca^2+^ release is taken up by mitochondria via MCU complex (Ahumada-Castro, Puebla-Huerta et al., 2021; Tanwar et al., 2021). However, the functional relevance of mitochondrial Ca^2+^ uptake in regulating melanogenesis remains totally unappreciated. To the best of our knowledge, this is the first study that demonstrates that mitochondrial Ca^2+^ uptake is a critical determinant of melanogenesis.

Mitochondrial Ca^2+^ signaling is emerging as a critical regulator of cellular physiology. However, its functional significance in skin biology in general and skin pigmentation in particular remains completely unappreciated. Here, we report a novel role of mitochondrial Ca^2+^ signaling in pigmentation biology. Using two independent cellular models and two distinct *in vivo* systems, we demonstrate a critical role of mitochondrial Ca^2+^ uptake in melanogenesis. Our data from both mouse LD pigmentation model and primary human melanocytes suggests that the mitochondrial Ca^2+^ uptake is directly proportional to the extent of pigmentation (**Figure 1**). Importantly, silencing and overexpression of MCU leads to abrogation and augmentation of melanogenesis respectively (**Figure 2**). Further, knockdown of negatively regulators of mitochondrial Ca^2+^ influx results in increase in melanogenesis (**Figure 3**). Moreover, we validated a critical role of MCU in regulating melanogenesis *in vivo* by performing MCU silencing; overexpression of human MCU and rescue studies in zebrafish (**Figure 4**). Additionally, MCU knock in mice shows enhanced epidermal pigmentation. Collectively, all this data establishes MCU mediated mitochondrial Ca^2+^ influx as a novel positive regulator of vertebrate pigmentation.

In order to characterize the molecular choreography working downstream of mitochondrial Ca^2+^ dynamics for regulating melanogenesis, we performed RNA seq on MCU and MCUb silenced B16 cells (**Figure 5**). Our unbiased RNA seq analysis led to identification of keratin 5, 7 and 8 as the bridges that connect mitochondrial Ca^2+^ signaling to melanosome biogenesis and maturation (**Figure 6 and 7**). So far, keratins are believed to regulate pigmentation via their influence on keratinocytes (Betz et al., 2006) and the role of keratin filaments in melanocytes remains largely unexplored. Although an earlier study had reported presence of intermediate filaments in melanocytes and suggested that these filaments can contribute to melanosome movement within melanocytes and melanosome transfer to keratinocytes (Jimbow & Fitzpatrick, 1975), their role in regulating melanosome biogenesis and maturation remained uncharacterized. To best of our understanding, this is the first report on role of melanocytic keratins in melanogenesis via their control over melanosome biogenesis and maturation (**Figure 7D-G**). Interestingly, a recent study demonstrated that in highly pigmented skin xenografts keratin 5 expression is higher than that in lightly pigmented vitiligo-like xenografts (Cario et al., 2020). The keratin 5 overexpression in highly pigmented xenografts was associated with increase in MITF (Microphthalmia-associated transcription factor) levels thereby suggesting that keratin 5 induces pigmentation via MITF. It is important to note that MITF is the master pigmentation regulator that plays an essential role in melanin synthesis, melanosome biogenesis and maturation (Nguyen & Fisher, 2019). Therefore, downstream of mitochondrial Ca^2+^ signaling, keratin 5 at least partially regulates pigmentation via MITF. However, further studies are needed to comprehensively characterize the role of keratins in melanosome biogenesis and maturation.

One of the intriguing findings of our study is that MCU silencing leads to NFAT2 nuclear translocation leading to enhanced keratin 5 expression (**Figure 8**), which in turn augments mitochondrial Ca^2+^ uptake (**Figure 6J-L**). Further, keratins act as positive regulators of melanogenesis by contributing to melanosome biogenesis and maturation. Therefore, this signaling module acts as a classical negative feedback loop that sustains both mitochondrial Ca^2+^ homeostasis and melanogenesis (**Figure 9D**). Given the central role of mitochondrial Ca^2+^ signaling and intermediate filaments in cellular physiology, this signaling module could be operational in a variety of cell types. Thereby, it might play an important role in several distinct physiological phenomena and pathological conditions. Further studies focused on understanding the relevance of crosstalk between mitochondrial Ca^2+^ signaling and intermediate filaments in different cellular models would establish broader significance of our work in human physiology.

The recent literature suggests that keratin filaments can regulate mitochondrial morphology, motility and respiration (Silvander et al., 2017; Steen et al., 2020). However, role of keratins in regulating mitochondrial Ca^2+^ uptake is not reported yet. We here show that keratin 5 positively regulate mitochondrial Ca^2+^ uptake. One of the members of intermediate filament family, desmin (a key muscle specific intermediate filament), is reported to regulate mitochondrial Ca^2+^ uptake in cardiac myocytes and skeletal muscle cells (Kostareva, Sjoberg et al., 2008; Smolina, Bruton et al., 2014). It is shown that aggregation prone desmin mutant decreases mitochondrial Ca^2+^ uptake and enhances cytosolic Ca^2+^ levels (Smolina et al., 2014). Interestingly, the mutant desmin did not perturb ER Ca^2+^ release thereby suggesting that it modulates mitochondrial Ca^2+^ uptake most likely by altering ER-mitochondrial crosstalk at Mitochondria-associated membranes (MAMs) (Schwarz & Leube, 2016; Sharma, Arora et al., 2020; Smolina et al., 2014). Independently, mitochondrial proteomics with desmin null and wild type mice hearts suggest that desmin controls VDAC expression (Fountoulakis, Soumaka et al., 2005). This can at least partially explain desmin mediated regulation of mitochondrial Ca^2+^ uptake. But the mechanism that links keratin 5 and mitochondrial Ca^2+^ uptake remains to be characterized. Future studies aimed at understanding the role of keratin 5 in regulating MAMs organization and expression/activity of mitochondrial Ca^2+^ uptake machinery will shed light on the underlying mechanism.

Another interesting observation of our study is clumping of melanosomes upon keratin 5 silencing in B16 cells (**Figure 7F**). Notably, aberrant distribution and clustering of cellular organelles is reported in skin cells of patients with keratin mutations (Uttam, Hutton et al., 1996). This suggests that our B16 cellular pigmentation model closely recapitulates the physiological pigmentation phenomenon. Moreover, these observations highlight the potential of targeting mitochondrial Ca^2+^ signaling for managing pigmentary disorders. Indeed, we demonstrate that inhibition of MCU with the FDA approved drug (mitoxantrone) leads to a significant decrease in αMSH stimulated physiological melanogenesis (**Figure 9A-B**). This suggests that MCU complex can be targeted for clinical management of pigmentary disorders especially the ones associated with keratin mutations and/or altered keratin expression. Keratin 5 mutations and altered expression are associated with a variety of pigmentary skin disorders such as Dowling Degos disease, epidermal bullosa with mottled pigmentation and senile lentigo (Betz et al., 2006, Cario et al.; 2020, Moog et al., 1999). Further, keratin 5 expression is directly correlated with the human pigmentary disorders i.e. very low in vitiligo (a depigmenting disorder) and extremely high in hyper-pigmentary disorders such as senile lentigo, photo exposed skin and Dowling Degos disease (Cario et al., 2020). Going forward, it would be worth to investigate expression and activity of MCU complex in different types of pigmentary disorders. Moreover, it is important to highlight that a variety of FDA approved drugs, which can modulate MCU complex activity have been characterized (Vishnu, Wilson et al., 2021). Therefore, future studies aimed at investigating the efficacy of these drugs in regulating pigmentation and thereby alleviating pigmentary disorders as well as aging associated hyperpigmentation are warranted.

Taken together, this study demonstrate an important role of mitochondrial Ca^2+^ signaling in melanogenesis, identify the molecular mechanisms that connect mitochondrial Ca^2+^ dynamics to melanosome biogenesis/maturation and report the potential of targeting mitochondrial Ca^2+^ influx for managing pigmentation. Furthermore, we show a novel MCU-NFAT2-Keratin 5 mediated feedback loop that maintains mitochondrial Ca^2+^ homeostasis and ensures optimal melanogenesis.

## Acknowledgements

This work was supported by the DBT/Wellcome Trust India Alliance Fellowship (IA/I/19/2/504651) awarded to Rajender K Motiani. Authors also acknowledge RCB core Institutional funding. M.M. is supported by the National Institutes of Health (R35GM145294, R01GM109882, R01HL142673 and R01DK135179). The authors thank members of the Motiani laboratory and Mohamed Trebak (University of Pittsburgh) for discussions and critical reading of the manuscript. The authors acknowledge Rajesh S Gokhale, DBT for insightful scientific discussions and Anurag Agrawal, Ashoka University for all the support. K.A. acknowledges her Junior Research Fellowship from DBT, India. The technical assistance of Ms. Nutan Sharma, Ms. Anushka Agrawal, Ms. Jaya Bharti Singh and Mr. Shyamveer is highly appreciated.

## Authors Contributions

Jyoti Tanwar: Methodology, Investigation, Visualization, Formal analysis, Writing Original draft preparation. Kriti Ahuja: Investigation, Formal analysis, Visualization. Akshay Sharma: Methodology, Investigation, Formal analysis, Visualization. Paras Sehgal: Investigation, Formal analysis. Gyan Ranjan: Investigation, Formal analysis. Farina Sultan: Formal analysis, Validation. Anshu Priya: Investigation, Formal analysis, Visualization. Manigandan Venkatesan: Investigation. Vamsi K Yenamandra: Resources. Archana Singh: Formal analysis, Visualization. Muniswamy Madesh: Resources. Sridhar Sivasubbu: Resources. Rajender K Motiani: Conceptualization, Supervision, Writing Original draft preparation, Reviewing and Editing, Project administration, Funding acquisition.

## Experimental Procedures

### Cell culture

B16-F10 cells were obtained from ATCC and cultured in DMEM (Sigma, Bangalore, India). Trypsin, Dulbecco‘s Phosphate Buffer Saline (DPBS), Versene, Fetal Bovine Serum (FBS) and additional cell culture grade reagents were obtained from Invitrogen, Waltham, MA, USA. Cells were cultured in DMEM medium supplemented with 10% Fetal Bovine Serum (heat inactivated) at 60–80% confluence and at 5% CO_2_ levels. Lightly and darkly pigmenting neonatal primary human melanocytes (LP and DP respectively) were procured from Invitrogen, Waltham, MA, USA. Cells were grown in Medium 254 supplemented with human melanocyte growth supplement-2 and maintained at 37 °C in a humidified incubator with 5% CO_2_ atmosphere. The maintenance and sub culturing of cells were carried out as per the manufacturer‘s instructions. Cells between passages 3–6 were used for experimentation.

### B16 Low-Density (LD) pigmentation-oscillator model

To set up the Low-Density (LD) pigmentation-oscillator model of B16s, cells were seeded at 100 cells/cm^2^ in DMEM supplemented with 10% Fetal Bovine Serum as described earlier (Motiani et al., 2018, Natarajan et al., 2014b).

### siRNA-based transfections

siRNA transfections were performed in T75 flasks on day 3 of the LD pigmentation model. 100 nM of siRNA (siRNAs from Dharmacon) was added per flask with a 1: 3 times V: V ratio of Dharmafect transfection reagent. siRNA and transfection reagent where mixed and incubated over the cells in OptiMEM (Gibco, Waltham, MA, USA) media for 4–6 h for achieving optimal transfection efficiency. Post transfection, the opti-MEM media was removed and then the day 3 culture media was added back to the cells.

siRNA transfections were performed in B16 high density cells using siRNAs from Dharmacon. Dharmafect was used for transfecting B16 cells. siRNA and transfection reagent where mixed and incubated over the cells in OptiMEM (Gibco, Waltham, MA, USA) media for 4–6 h for achieving optimal transfection efficiency. As reported earlier, siRNA transfection in primary melanocytes was done using Nucleofection Kit (Lonza, VPD-1003, U-024 program) (Sultan et al., 2022). 100 nM of siRNA (smartpool siRNAs from Dharmacon) and 0.5ug Pmax-gfp plasmid DNA was added per condition (7-10 lac cells). Media was changed after 24 hours of transfection. Cells were harvested post 72 hours of transfection to capture phenotypic and protein expression changes. The siRNAs were procured from Dharmacon.

The catalog numbers of siRNAs used in the study are included in Table 1.

**Table 1.** Details of siRNAs used.

| siRNA | Catalog Number |
| --- | --- |
| siNT | D-001810-10-20 |
| siMCU (Mouse) | J-062849-05-0010 |
| siMCU (Human) | L-015519-02-0010 |
| siMCUb (Mouse) | L-061742-01-0005 |
| siMCUb (Human) | L-016108-02-0010 |
| siMICU1 (Mouse) | L-053388-01-0005 |
| siKeratin 7 (Mouse) | L-063818-01-005 |
| siKeratin 8 (Mouse) | L-043866-01-005 |
| siKeratin 5 (Mouse) | L-048162-01-0005 |
| siKeratin 5 (Human) | L-011067-00-0005 |

### qRT-PCR analysis

For mRNA extraction cells were processed with Qiagen RNeasy kit (Catalog #74106). mRNA was then converted to cDNA using high-capacity cDNA reverse transcription kit from ThermoFisher (Waltham, MA, USA) (Catalog #4368814). Real-time PCR reactions were performed using SYBR green in Quant Studio 6 Flex from Applied Biosystems. The data were analyzed with Quant Studio real-time PCR software version 1.3. The expression of gene of interest was normalized to that of the housekeeping gene GAPDH. Primers were designed using Primer3 and checked by the NCBI Primer BLAST tool. Gene-specific primers were obtained from Eurofins.

**Table 2.**
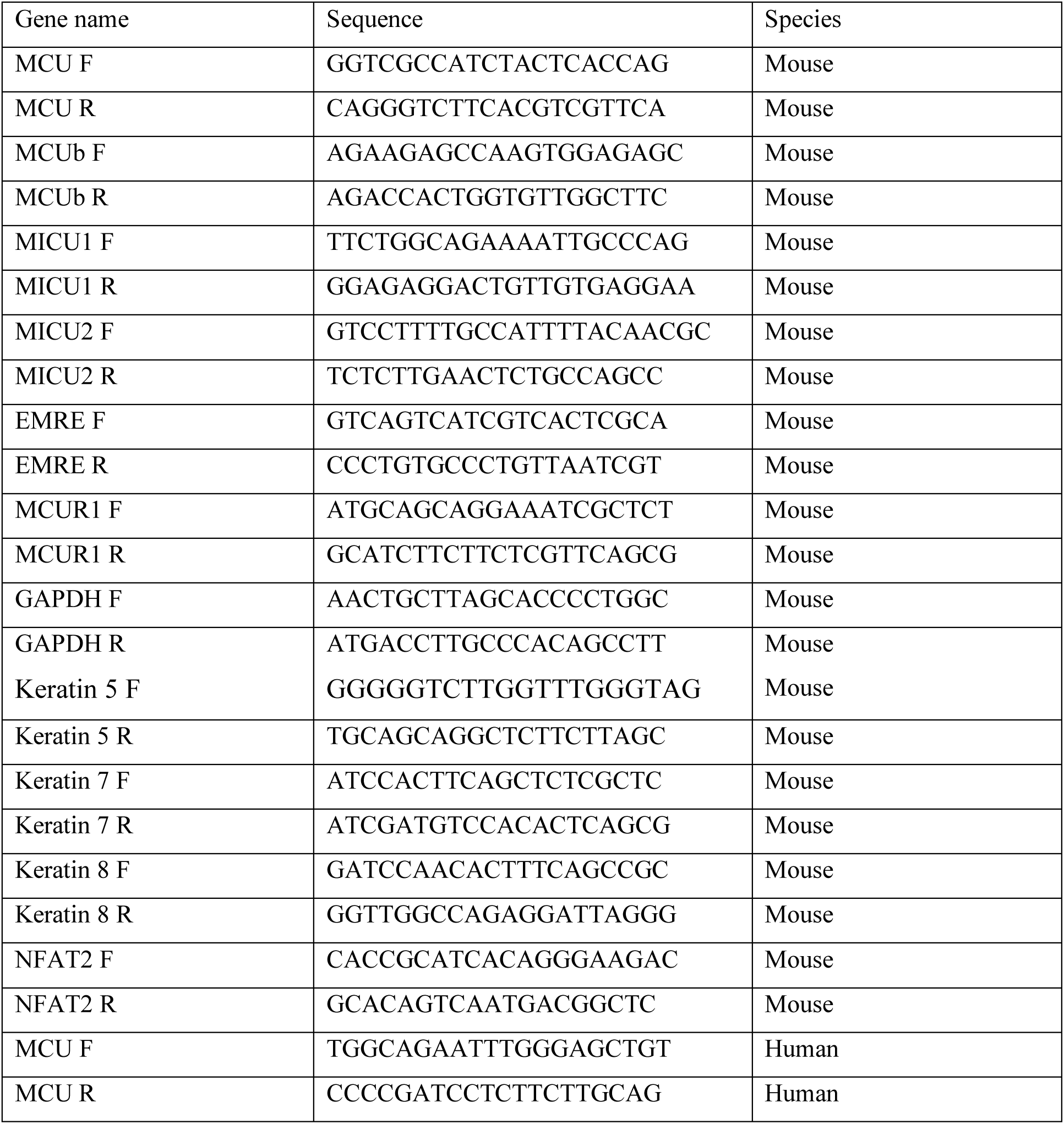
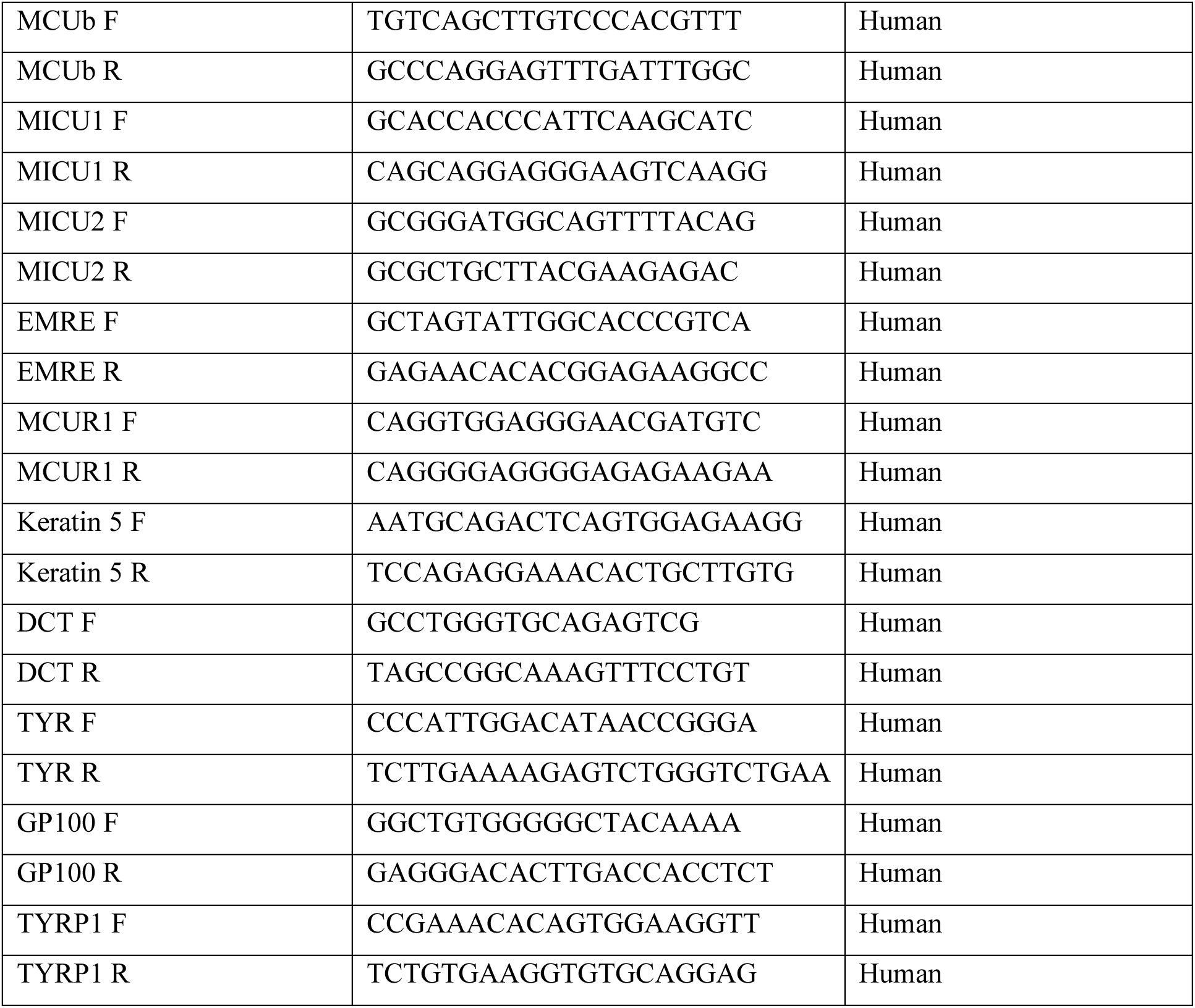
List of qRT-PCR primers (F stands for forward primer and R for reverse primer)

### Western blotting

Cells were lysed using NP40 lysis buffer supplemented with protease inhibitors. Typically, 50–100 µg proteins were subjected to SDS-PAGE (7.5–10%). Proteins from gels were then electro-transferred onto PVDF membranes. After blocking with 5% non-fat dry milk (NFDM) dissolved in Tris-buffered saline containing 0.1% Tween 20 (TTBS), blots were probed overnight at 4 °C, with specific primary antibodies in TTBS containing 2% NFDM. The primary antibodies used were typically procured from Abcam, Cell signaling and were used at 1:500–1:2000 dilutions. The following day, membranes were incubated for 2 h at room temperature with a horseradish-peroxidase-conjugated anti-mouse or anti-rabbit IgG antibody in TTBS containing 2% NFDM. Detection was performed using the enhanced chemiluminescence reagent (ECL Western blotting detection reagents; Amersham Biosciences). Quantification of bands was performed by densitometry using the ImageJ software. The catalogue number and company name for the antibodies are provided in Table 3.

**Table 3.**
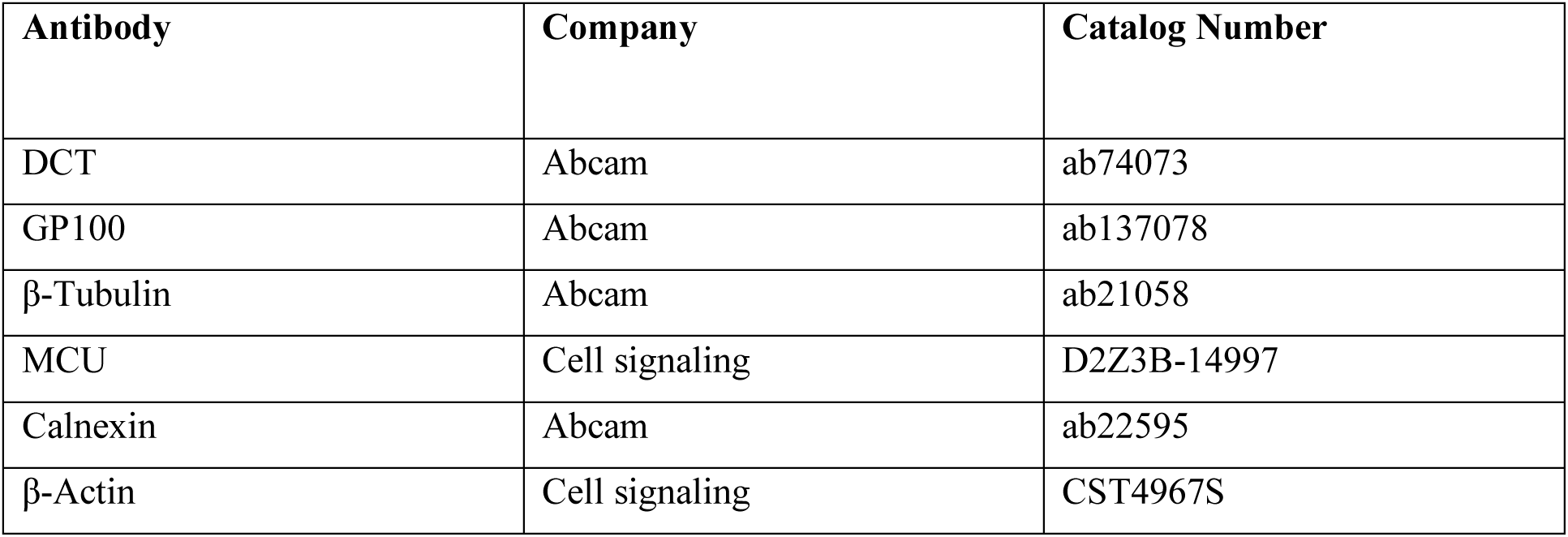
Details of antibodies used.

| Antibody | Company | Catalog Number |
| --- | --- | --- |
| DCT | Abcam | ab74073 |
| GP100 | Abcam | ab137078 |
| β-Tubulin | Abcam | ab21058 |
| MCU | Cell signaling | D2Z3B-14997 |
| Calnexin | Abcam | ab22595 |
| β-Actin | Cell signaling | CST4967S |

### Tyrosinase activity/DOPA assay

Tyrosinase-enzyme activity was checked in cell lysates by performing DOPA assay as previously reported (Fuller, Drake et al., 2000). Briefly, cell lysates were prepared in NP-40 lysis buffer and an equal amount of protein was run on a gel under nonreducing/native conditions. The gel was then immersed in phosphate buffer supplemented with tyrosinase substrate L DOPA (Sigma Chemicals, Bangalore, India). Enzyme activity corresponded to the formation of black-color pigment.

### Melanin content assay

Melanin-content assay was performed as described earlier (Kageyama, Oka et al., 2004). The cells were lysed in 1N NaOH by heating at 80°C for 2 hours and then absorbance was measured at 405nm. Melanin content was estimated by interpolating the sample readings on the melanin standard curve (µg/mL) obtained with synthetic melanin. Zebrafish embryos were lysed in 1N NaOH by heating at 90°C for 6 hours.

### Mitochondrial calcium imaging

For performing mitochondrial Ca^2+^ imaging, B16 cells (0.5 x10^6^ cells/well) were plated on confocal dishes (SPL Life Sciences, Korea). After 24hours post B16 cell plating (at 60% confluency), pCMV CEPIA2mt (Addgene plasmid # 58218) plasmid (1.5 µg) was overexpressed using Lipofectamine 2000 (Invitrogen, 11668-019). After 24 hours post transfection, cells were washed 3 times and bathed in HEPES-buffered saline solution (140 mM NaCl, 1.13 mM MgCl2, 4.7 mM KCl, 2 mM CaCl2, 10 mM D-glucose, and 10 mM HEPES; pH 7.4) for 5 min before Ca^2+^ measurements. Nikon Eclipse Ti2 microscope was used to record and analyze fluorescence images of several cells using 60X oil objective. For CEPIA2mt following excitation/emission wavelengths: (488 nm/500–550 nm) were used. 100µM histamine in Ca^2+^ free bath solution was used to release Ca^2+^ from the ER via IP3Rs. Experiments were performed at least 3 times and the final data are plotted in the form of violin plots where the number of cells is denoted as “n”. For measuring mitochondrial Ca^2+^ on different days in Low-Density (LD) pigmentation-oscillator model of B16s, cells were seeded at 100 cells/cm^2^ on confocal dishes. pCMV CEPIA2mt plasmid (1.5 µg) was overexpressed using Lipofectamine 2000, 24hours before measuring mitochondrial Ca^2+^. pCMV CEPIA2mt was a gift from Masamitsu lino (Addgene plasmid # 58218).

For measuring mitochondrial Ca^2+^ in lightly pigmented (LP) and and darkly pigmented (DP) primary human melanocytes using Rhod-2 AM (543 nm/580–650 nm) dye, primary human melanocytes (1 x10^6^ cells/dish) were plated on confocal dishes. After 24hours post cell plating (at 50-60% confluency), cells were washed with media without FBS and antibiotic antimycotic agents. Further, cells were incubated in media containing 1 µM Rhod-2 AM at 37°C for 30 min. Cells were washed and kept in HBSS (HEPES-buffered saline solution (140 mM NaCl, 1.13 mM MgCl_2_, 4.7 mM KCl, 2 mM CaCl_2_, 10 mM D-glucose, and 10 mM HEPES, adjusted to pH 7.4 with NaOH)) containing 5 mM CaCl_2_ for imaging. The cells were stimulated with 100µM histamine in HBSS containing 5 mM CaCl_2_. Nikon Eclipse Ti2 microscope was used to record and analyze fluorescence images of several cells using 60X oil objective. The data was acquired every 1s intervals for 5–10 min.

### MCU overexpression in B16 cells

B16 cells were seeded 24hrs before transfection at a density of 1.0 x106 cells/well in 6 well plates. Human pDEST47-MCU-GFP plasmid (1.5 µg) was overexpressed in B16 cells plated at 60% confluency using Lipofectamine 2000 (Invitrogen, 11668-019). After 18 hours post transfection, 1µM α-MSH treatment (Sigma-Aldrich, M4135) was given for 30 hours followed by cell termination. Human MCU-GFP plasmid was a gift from Vamsi Mootha (Addgene plasmid # 31732).

### Mitoxantrone treatment in *α*MSH-induced pigmentation assay

In B16 cells, seeded at HD, pigmentation was induced by adding 1µM αMSH (Sigma Aldrich, M4135) for 48 h. Cells were pretreated (for 1 hour) with 1µM Mitoxantrone (Sigma Aldrich, M6545) followed by addition of 1µM αMSH for 48 h. After 48 h cells were terminated and pellets were made. Further, mean pixel intensity of pellets was calculated using ImageJ software.

### Generation of KRT5 overexpression construct

Human KRT5 overexpression plasmid was generated by sub cloning human KRT5 cDNA into mCherryC1 vector at EcoRI/XhoI sites. KRT5 cDNA was PCR amplified using pBabe mRFP1-KRT5 (Addgene#59493) as template using Phusion High Fidelity Polyemrase (F503, Thermo). The amplified PCR product was restriction digested using EcoRI/XhoI (NEB). Simultaneously, mCherryC1 empty vector was also digested along with shrimp alkaline phosphatase (NEB) treatment. This was followed by gel purification of digested vector and PCR product. Vector and insert ligation was performed using Rapid DNA ligation Kit (K1422, Thermo). The ligation mix was transformed into chemically competent E.coli DH5α cells followed by plating onto Kanamycin containing agar plates. Positive clones were screened by restriction digestion analysis of plasmid DNA obtained by miniprep. Primers utilized for cloning are as follows: hKRT5CDS FP 5’-AAACTCGAGCCATGTCTCGCCAGTCAA-3’ hKRT5CDS RP 5’-CCCGAATTCTTAGCTCTTGAAGCTCTTCC-3’

### Cloning of Keartin5 promoter

Mouse Keratin5 promoter sequence was obtained from Eukaryotic Promoter Database (EPD), which was followed by NCBI-BLAST analysis of the sequence to identify mRNA start site and first codon. Primers were designed to amplify an 1106bp region (-1000 to + 106, w.r.t. to start codon) of the KRT5 promoter. KRT5 core promoter -1000 to +106 (KRT5PWT) was amplified from B16F10 genomic DNA, isolated using DNeasy Blood and Tissue Kit (69504, Qiagen) as per manufacturer’s protocol. This was followed by PCR amplification of the target region using Phusion High Fidelity Polymerase (F503, Thermo), which was further cloned into pGL4.23 luciferase reporter vector (Promega) at the KpnI/HindIII sites. Cloned promoter was verified by restriction digestion and sequencing to verify its identity. Primers utilized for cloning are as follows: KRT5P FP 5’-TTAAGGTACCGTGTTTGCGGGCGG-3’ KRT5P RP 5’-GGTTAAGCTTTGGCGAGACATGATGG-3’

### In vitro luciferase assay

B16 cells were seeded 24hrs before transfection at a density of 0.5x10^5^ cells/well in 24 well plates. Cells were transfected with KRT5PWT along with eGFPc1-huNFATc1EE-WT (Addgene#24219) as indicated, using Turbofect transfection reagent (R0532, Thermo) as per manufacturer’s protocol. Renilla luciferase control plasmid was utilized for transfection normalization in all experiments. 48h post transfection, cells were assayed for luciferase activity using the dual luciferase assay kit (E1910, Promega, Madison, WI) as per manufacturer’s protocol. Data is representative of 3 biological replicates with 3 technical replicates each.

### Transmission Electron Microscopy

Transmission electron microscopy was performed on siNT, siMCUb and siMCUb+Keratin 5 B16 cells on LD day 6 using standard protocols. Briefly, cells were fixed overnight in fixative containing 2.5% glutaraldehyde and 4% paraformaldehyde, gradually dehydrated in graded series of ethanol and embedded in Epon 812 resin. Ultrathin sections were cut and stained with uranyl acetate and lead citrate and images were captured using a transmission electron microscope (JEM-1400Flash).

### Zebrafish husbandry

Zebrafish used in this study were housed in CSIR-Institute of Genomics and Integrative Biology, India with proper standard ethical protocols approved by the Institutional Animal Ethics Committee of CSIR-Institute of Genomics and Integrative Biology, India with care to minimize animal suffering.

### Morpholino design and microinjections

Antisense morpholino (MO) oligonucleotides were designed against MCU of zebrafish to block the translation and were ordered from Gene Tools, USA. The MO were reconstituted in nuclease free water (Ambion, USA) according to the recommended protocol by GENE tools to get 1mM concentration and was stored in -20 °C for further experiments. MO was injected at a single cell stage and embryos were cultured in E3 water at 28°C. Dose titration for MO was done using 3 different concentrations (100, 200, 500 uM) and the optimal dose concentration 100uM was selected based on survival and phenotype percentage. Scrambled MO was used as a negative control for the experiments. The embryos were screened for pigmentation phenotype at 30 hpf, 36hpf and 48 hpf. Imaging was done using Nikon microscope. The sequence of morpholino used is provided in Table 4.

**Table 4.**
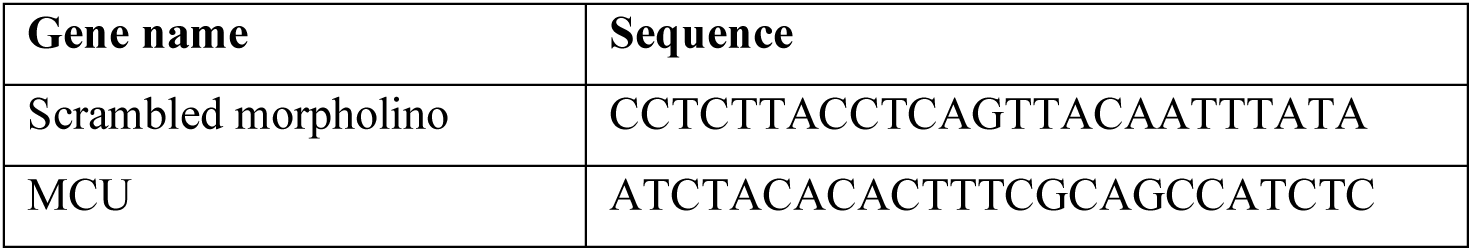
Details of morpholino used.

### Overexpression and complementation/rescue assay of MCU in zebrafish

Plasmid containing CDS sequences of human MCU were taken from Addgene and restriction digestion was performed. In vitro transcription (IVT) was performed using T7 mMassage mMachine kit (invitrogen) and purified using RNA purification column (Roche). MCU-IVT was injected at single cell stage and embryos were cultured in E3 water. Dose titration with different concentration of IVT (50, 100, 200 ng/ul) was performed for the overexpression experiment. The optimal concentration 50ng/ul was selected based on survival and phenotype percentage of the embryos. The embryos with overexpression were screened for pigmentation phenotype at 24 hpf and 48 hpf.

Complementation/rescue assays were performed in zebrafish using MCU knockdown embryos. A cocktail of MCU MO (100uM) and Human MCU IVT (50ng/ul) was injected into single cell stage zebrafish. RFP IVT was used as a negative control for the complementation assay experiment. The embryos were screened for pigmentation phenotype at 24 hpf and 48 hpf to understand the level of rescue in complementation assay. Imaging was done using Nikon microscope.

### MCU KI mice

The hypermorphic MCU-C96A-knockin (KI) mice was generated and reported in our earlier study (Tomar et al., 2019). MCU-C96A corresponds to human MCU C97A and results in hyperactivation of MCU channel activity (Tomar et al., 2019). In order to phenotypically assess the role of MCU in mice epidermal pigmentation, we took tail pictures of MCU KI mice and corresponding wild type control mice with a digital camera. To quantitate the differences in the epidermal pigmentation between WT mice and MCU KI mice, we calculated mean pixel intensity of mice tail using ImageJ software. We analyzed 30 data points from 3 independent tails of wild type mice and MCU KI mice (10 data points/mice and 3 mice/condition).

### RNA sequencing

LD day 6 samples siNT, siMCU and siMCUB were sent for mRNA sequencing to Clevergene, Bengaluru, India. The sequence data was generated using Illumina NovaSeq. Data quality was checked using FastQC and MultiQC software. GRCm39 was used as the reference genome. Gene level expression values were obtained as read counts using feature counts software and normalized counts were obtained. Fold change was calculated for siMCU and siMCUb samples with respect to non-targeting siRNA control. Genes showing log2 fold change more than +1 in expression with respect to siNT control were taken as upregulated genes in siMCU and siMCUb. While a cut off of log2 fold change less than -1 was set for determining downregulated genes in siMCU and siMCUb. These upregulated and downregulated genes were analyzed for pathway enrichment using DAVID software. Pathway enrichment analysis was performed based on the Gene Ontology (GO) database. Enriched pathways were plotted with enrichment scores in Graphpad. To determine oppositely regulated pathways in siMCU and siMCUb, common pathways up in siMCU and down in siMCUb and vice versa (down in siMCU and up in siMCUb) were plotted. Further, to get common but oppositely regulated genes, the Venn diagram was plotted for genes up in siMCU and down in siMCUb and vice versa.

### Measurement of NFAT2 nuclear translocation

To evaluate the effects of MCU mediated Ca^2+^ flux on NFAT2 nuclear translocation, B16 cells were seeded at a density of 20,000 cells/ well in a 6 well plate. Cells were then transfected with smart pool siRNA against mouse MCU (Dharmacon) or control siRNA (Dharmacon) at a concentration of 100nM using DharmaFect transfection reagent according to manufacturer’s protocol. 24hrs post transfection cells were reseeded into glass bottom imaging dishes. 72hrs post transfection cells from both siMCU and siNT conditions were transfected with 1ug of eGFP-NFAT2 overexpression plasmid (Addgene#24219) using Lipofectamine 2000 reagent according to manufacturer’s protocol. 96hrs post siRNA transfections live cell imaging was performed. Cells were washed thrice with Ca^2+^ containing HBSS and imaged in Ca^2+^ containing HBSS bath using a Nikon Eclipse Ti2 microscope at 40x magnification. eGFP-NFAT2 was excited using 488nm laser and emission was captured at 510nm. eGFP-NFAT2 translocation in response to 100uM histamine stimulation was analysed in real-time using the equation:

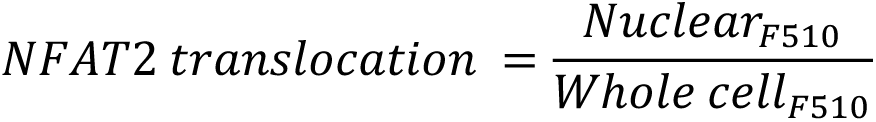

Further, confocal microscopy was also performed using a Carl Zeiss LSM 880 laser scanning confocal microscope at 63X (with oil) magnification at the indicated time points to demonstrate nuclear translocation of NFAT2 in B16 cells with knockdown of MCU in response to 100uM histamine stimulation.

### Statistical analysis

All statistical analysis was performed using GraphPad Prism 8 software. All experiments were performed at least 3 times. Data are presented as mean ± SEM. Either unpaired student’s t-test or One-sample t-test was performed for determining statistical significance between 2 experimental samples, whereas one-way ANOVA was performed for the comparison of 3 samples. A p-value < 0.05 was considered as significant and is presented as “*”; p-value < 0.01 is presented as “**”; p-value < 0.001 is presented as “***” and *p* < 0.0001 is presented as “****”.

---

**Supplementary Figure 1 supporting Main Figure 2.**
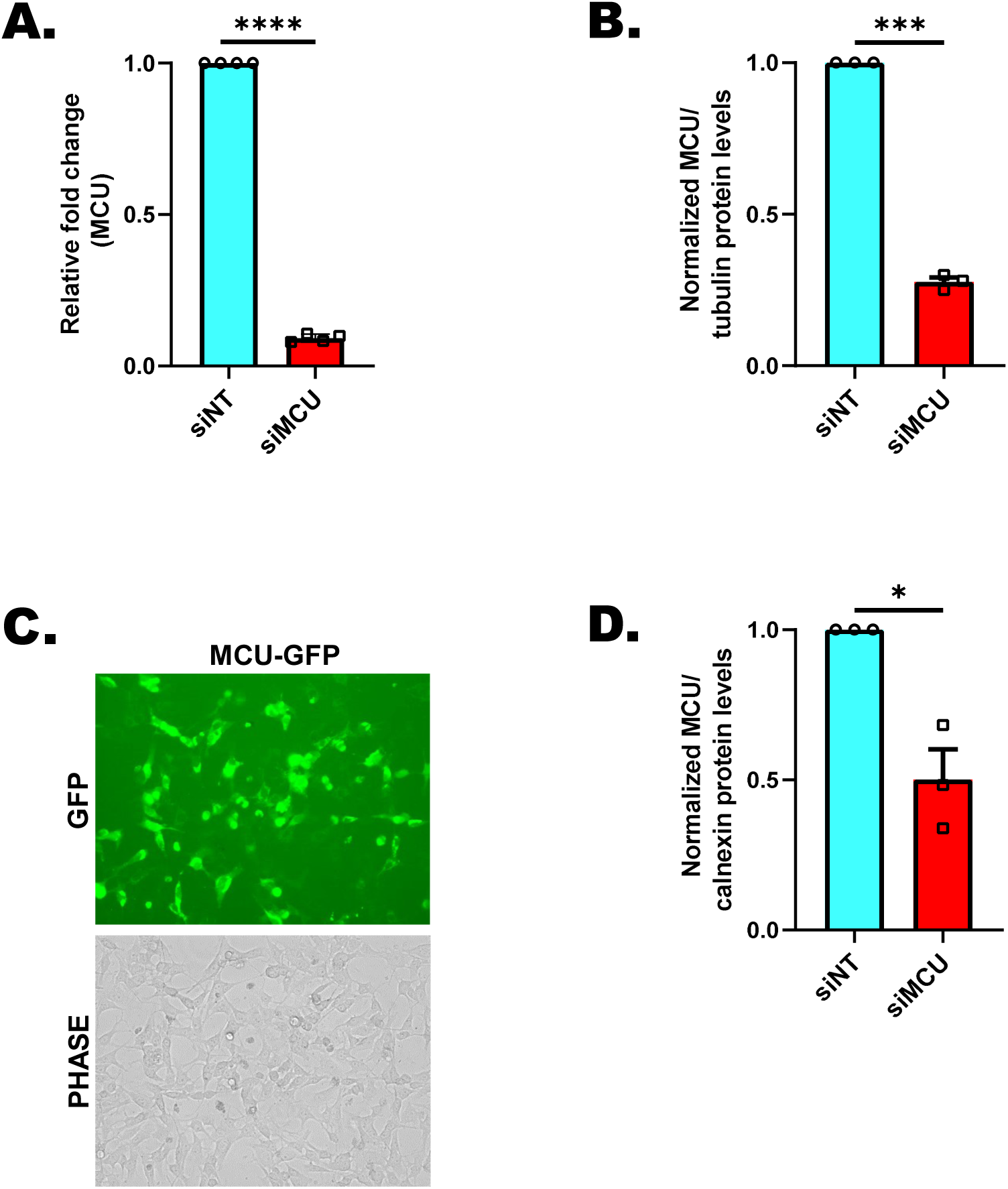
MCU positively regulates melanogenesis. **(A)** qRT–PCR analysis showing decrease in MCU mRNA expression upon MCU silencing in B16 cells (N=4). **(B)** Densitometric quantitation showing MCU levels on LD day 6 in siNT control and siMCU condition (N=3). **(C)** Representative GFP and bright field images showing MCU-GFP transfected B16 cells (Scale = 100µm) (N=3). **(D)** Densitometric quantitation showing MCU levels in siNT control and siMCU condition in primary human melanocytes (N=3). Data presented are mean ± S.E.M. For statistical analysis, one sample *t*-test was performed for panels A, B, D using GraphPad Prism software. * *p* <0.05; *** *p* < 0.001 and **** *p* < 0.0001.

**Supplementary Figure 2 supporting Main Figure 3.**
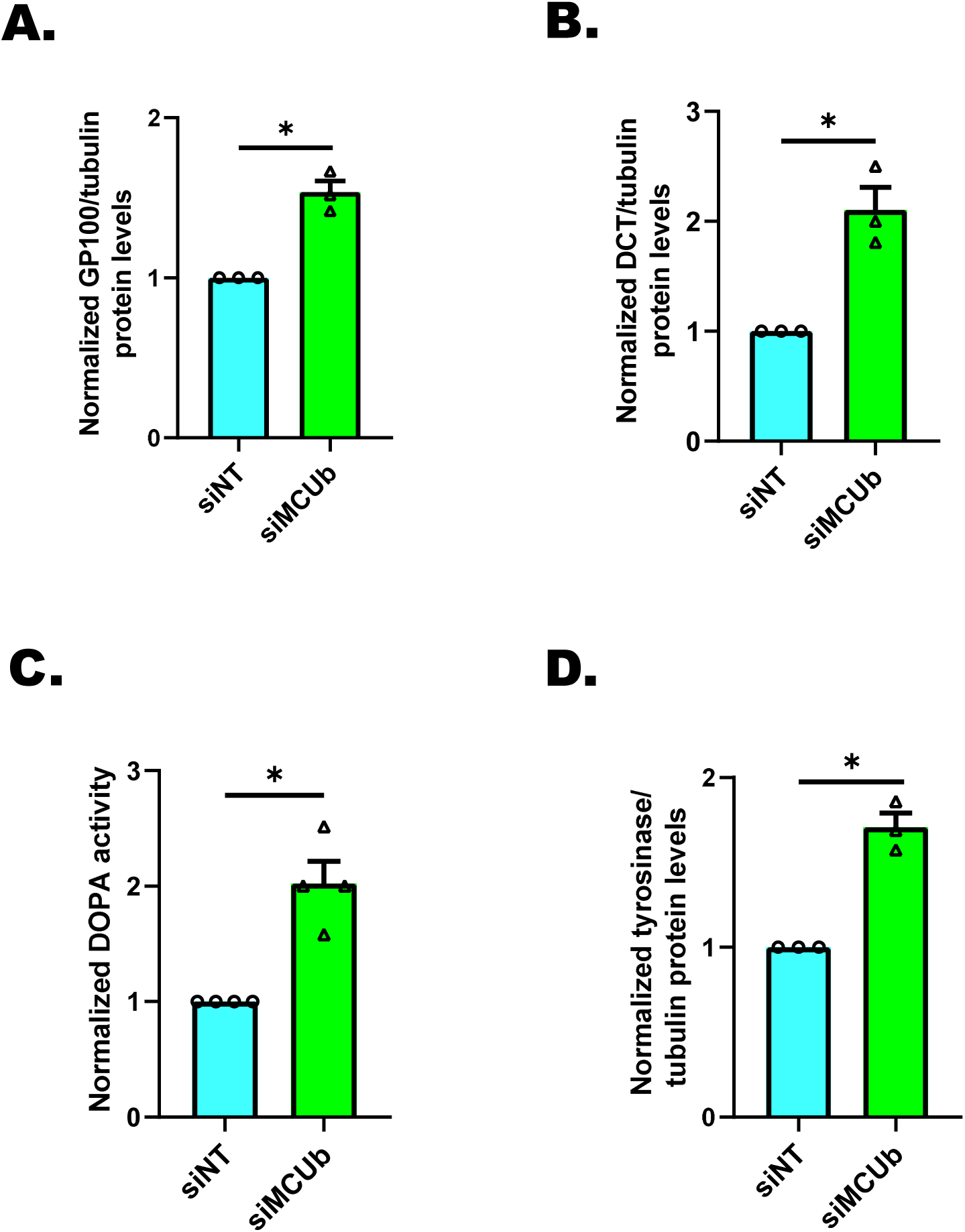
MCUb and MICU1 negatively regulate melanogenesis. **(A)** Densitometric quantitation showing GP100 levels on LD day 6 in siNT control and siMCUb condition (N=3). **(B)** Densitometric quantitation showing DCT levels on LD day 6 in siNT control and siMCUb condition (N=3). **(C)** Densitometric quantitation showing activity of tyrosinase enzyme on LD day 6 in siNT control and siMCUb condition (N=4). **(D)** Densitometric quantitation showing Tyrosinase levels on LD day 6 in siNT control and siMCUb condition (N=3). Data presented are mean ± S.E.M. For statistical analysis, one sample *t*-test was performed for panels A-D using GraphPad Prism software. Here * *p* <0.05.

**Supplementary Figure 3 supporting Main Figure 4.**
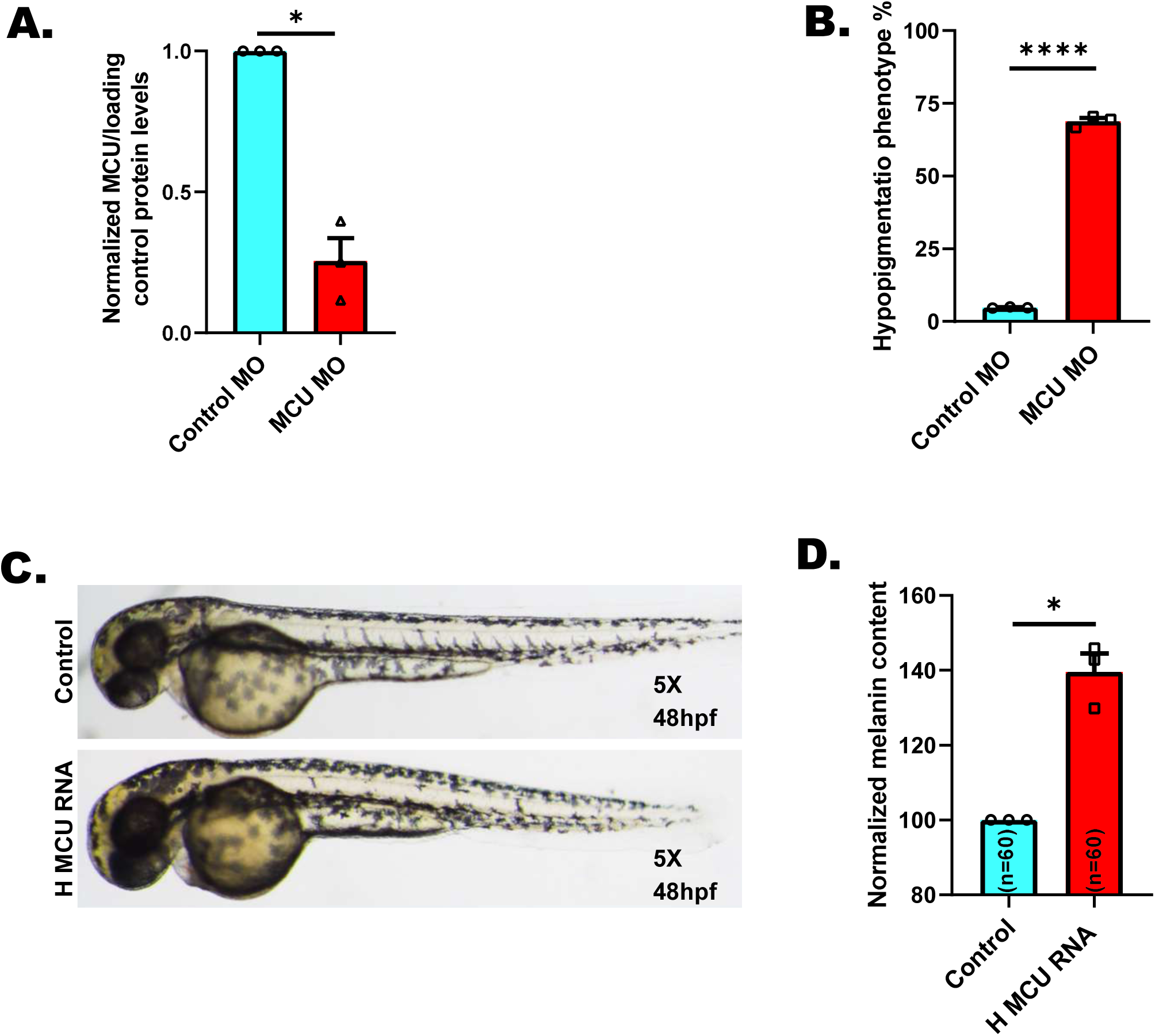
MCU regulates pigmentation *in vivo*. **(A)** Densitometric quantitation showing MCU levels in control MO and MCU MO (N=3). **(B)** The hypopigmentation phenotype analyzed at 30hpf in around 200 zebrafish embryos from three independent sets of injections (N=3 independent experiments with ∼200 embryos/condition). **(C)** Representative bright-field images of zebrafish embryos injected with either control or human MCU RNA at 48hpf (N=3 independent experiments with ∼200 embryos/condition). **(D)** Melanin-content estimation of zebrafish embryos injected with either control or human MCU RNA injection in 60 zebrafish embryos from three independent sets of injections (N=3 independent experiments with 60 embryos/condition). Data presented are mean ± S.E.M. For statistical analysis, one sample *t*-test was performed for panel A, D while unpaired *t*-test was performed for panel B using GraphPad Prism software. Here, * *p* <0.05 and **** *p* < 0.0001.

**Supplementary Figure 4 supporting Main Figure 6.**
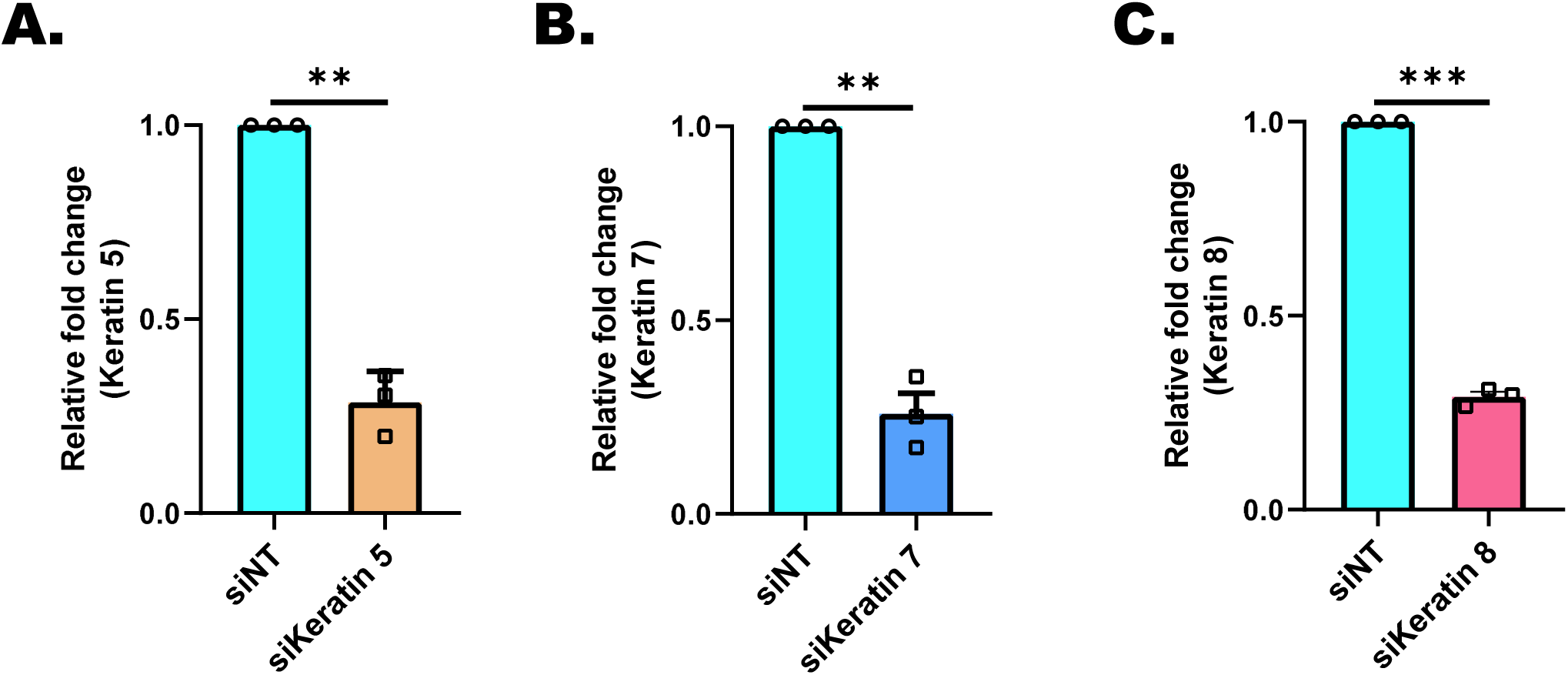
Keratins regulate melanogenesis. **(A)** qRT–PCR analysis showing decrease in keratin 5 mRNA expression upon keratin 5 silencing in B16 cells (N=3). **(B)** qRT–PCR analysis showing decrease in keratin 7 mRNA expression upon keratin 7 silencing in B16 cells (N=3). **(C)** qRT–PCR analysis showing decrease in keratin 8 mRNA expression upon keratin 8 silencing in B16 cells (N=3). Data presented are mean ± S.E.M. For statistical analysis, one sample *t*-test was performed for panels A-C using GraphPad Prism software. Here, ** *p* < 0.01 and *** *p* < 0.001

**Supplementary Figure 5 supporting Main Figure 8.**
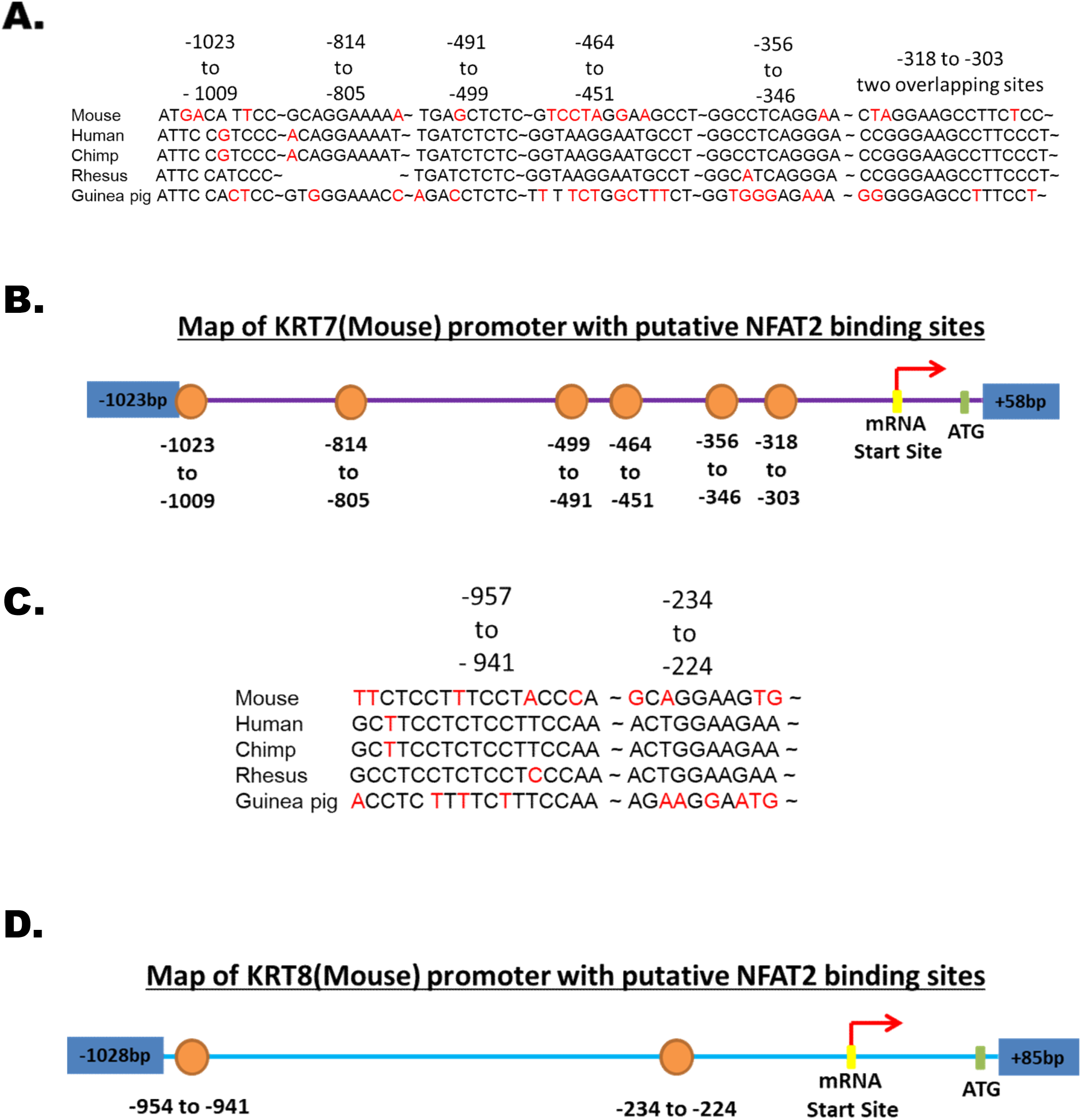
NFAT2 connects MCU to keratins expression. **(A)** Multispecies sequence alignment of putative NFAT2 binding sites in the mouse keratin 7 (KRT7) core promoter. **(B)** Schematic representation of putative NFAT2 binding sites in the mouse KRT7 core promoter. **(C)** Multispecies sequence alignment of putative NFAT2 binding sites in the mouse keratin 8 (KRT8) core promoter. **(D)** Schematic representation of putative NFAT2 binding sites in the mouse KRT8 core promoter.

